# Endothelium-released Dll4 drives sarcopenia and systemic aging

**DOI:** 10.64898/2026.11.25.769149

**Authors:** Shin Fujimaki, Naoki Horii, Ryosuke Okino, Huascar Pedro Ortuste Quiroga, Kodai Nakamura, Yuri Miura, Kota Sawada, Shoma Iritani, Yoko Mikumo, Yuki Goda, Sayoko Fujimura, Kei-ichiro Yasunaga, Shingo Usuki, Yukie Takahashi, Yuta Miyamoto, Takaichi Fukuda, Takahiro Eguchi, Naoki Ito, Takashi Minami, Naoto Kajitani, Minoru Takebayashi, Daisuke Ando, Yusuke Ono

**Affiliations:** Department of Muscle Development and Regeneration, Institute of Molecular Embryology and Genetics, Kumamoto University, Kumamoto, Japan; Research Team for Aging Science, Tokyo Metropolitan Institute for Geriatrics and Gerontology, Itabashi-ku, Tokyo, Japan; Research Team for Mechanism of Aging, Tokyo Metropolitan Institute for Geriatrics and Gerontology, Tokyo, Japan; Liaison Laboratory Research Promotion Center, Institute of Molecular Embryology and Genetics, Kumamoto University, Kumamoto, Japan; Department of Anatomy and Neurobiology, Graduate School of Medical Sciences, Kumamoto University, Kumamoto, Japan; Brain-Skeletal Muscle Connection in Aging Project Team, Geroscience Research Center, National Center for Geriatrics and Gerontology, Aichi, Japan; Division of Molecular and Vascular Biology, IRDA, Kumamoto University, Kumamoto, Japan; Department of Psychiatry and Neuroscience, Center for Metabolic Regulation of Healthy Aging, Faculty of Life Sciences, Kumamoto University, Kumamoto, Japan; Faculty of Education, Graduate Faculty of Interdisciplinary Research, University of Yamanashi, Kofu, Yamanashi, Japan

## Abstract

Lifelong preservation of skeletal muscle function is essential for healthy aging. Sarcopenia, the age-related loss of muscle mass and strength, arises from aging itself (primary sarcopenia) or chronic diseases (secondary sarcopenia). Although these forms frequently coexist, whether they share a therapeutically targetable mechanism remains unclear. Here we identify soluble, full-length Delta-like ligand 4 (Dll4), released from endothelial cells, as a common driver of both forms of sarcopenia. Endothelium-derived Dll4 increases with age and accumulates in interstitial spaces through binding to type I collagen, where it selectively activates Notch2. Chronic Dll4 exposure impairs muscle function, induces endothelial senescence, and promotes vascular regression. Conversely, sustained Dll4 inhibition attenuates primary sarcopenia, preserves vascular integrity, ameliorates systemic aging phenotypes, and extends lifespan, even when initiated in aged mice. Studies in human cohorts and mouse models further uncover a kidney–vascular–muscle axis linking renal dysfunction to elevated circulating Dll4, whose blockade prevents secondary sarcopenia. Together, these findings indicate that endothelium-released Dll4 acts as a shared mediator of primary and secondary sarcopenia and systemic aging, highlighting its inhibition as a promising strategy to extend healthspan and lifespan.

## Introduction

Aging is a systemic and local process characterized by a progressive decline in tissue homeostasis and physiological resilience, ultimately increasing susceptibility to frailty, chronic diseases, organ dysfunction, and mortality (1). Sarcopenia, the age-related loss of muscle mass and strength, is a major determinant of physical function and healthspan in older adults (2).

Age-related sarcopenia (primary sarcopenia) is attributed to physiological muscle aging in the absence of other identifiable causes. Muscle aging involves multiple structural, cellular, and molecular alterations, including myofiber atrophy, changes in muscle fiber-type composition, neuromuscular junction (NMJ) degeneration, a decline in satellite cell numbers, chronic low-grade inflammation, impaired proteostasis, and reduced responsiveness to anabolic stimuli (3, 4). Secondary sarcopenia results from factors other than aging (including physical inactivity, malnutrition, and systemic diseases such as chronic kidney disease and metabolic disorders). Primary and secondary sarcopenia are not mutually exclusive but frequently coexist, particularly in older adults with multiple comorbidities (2). Therefore, identifying and targeting shared molecular and cellular mechanisms underlying both conditions may constitute a promising therapeutic strategy to preserve muscle function and extend healthspan.

Muscle mass and function are regulated not only by intrinsic processes within myofibers but also by the surrounding tissue microenvironment, which comprises diverse cell populations, including mesenchymal stromal cells, immune cells, and endothelial cells (ECs) (3). Although primary and secondary sarcopenia have distinct etiological factors, they may converge on shared pathological processes within the microenvironment (2, 5). However, despite growing recognition of the importance of microenvironment, it remains unclear how intercellular communication contributes to sarcopenia (6).

The Notch signaling pathway is a highly conserved intercellular communication system that plays a pivotal role in regulating diverse cellular functions and maintaining tissue homeostasis through direct cell–cell interactions (7). In mammals, four Notch receptors (Notch1–4) and five canonical ligands belonging to the Delta/Serrate/LAG-2 family have been identified: Jagged1, Jagged2, and Delta-like ligands 1, 3, and 4 (Dll1, Dll3, and Dll4) (8). Notch signaling is crucial for regulating satellite cell function in adult skeletal muscle (9–18). We previously demonstrated that Notch2 is expressed not only in satellite cells but also in differentiated multinucleated myofibers, and that EC-derived Dll4 activates myofiber Notch2, thereby promoting muscle atrophy under conditions of physical inactivity and diabetes (9, 19). However, whether the endothelial Dll4–myofiber Notch2 signaling axis contributes to the development of sarcopenia remains unknown. Here, we investigated the role of this signaling axis in skeletal muscle aging.

## Results

### Dll4–Notch2 axis inhibition ameliorates sarcopenia

To determine the role of Notch2 in muscle aging, we generated myofiber-specific Notch2-deficient mice by crossing *Notch2^f/f^* (N2^f/f^) mice (20) with myosin light chain 1f promoter-driven Cre recombinase-expressing (*Mlc1f^Cre/+^*) mice (21) (*Mlc1f^Cre/+^*;*N2^f/f^,* N2^−/−^), as previously described (19). N2^−/−^ mice were born at the expected Mendelian frequency, exhibited normal growth, and showed no overt abnormalities in skeletal muscle during young adulthood. However, aging phenotypes in muscle were markedly attenuated in N2^−/−^ mice. Control mice displayed the expected age-associated reductions in muscle mass, while aged N2^−/−^ mice were largely protected from these declines (Fig. 1A,B). Consistent with these findings, maximal muscle force generation and grip strength were substantially preserved in aged N2^−/−^ mice (Fig. 1C–E). To examine molecular pathways downstream of Notch2, we performed unbiased bulk RNA-sequencing (RNA-seq) of aged muscle, and Kyoto Encyclopedia of Genes and Genomes (KEGG) pathway analysis (22) revealed age-associated upregulation of proteasome-related pathways in control muscle, consistent with a previous study (23). This upregulation was largely absent in N2*^−/−^*mice (Fig. 1F), as further validated by quantitative PCR (qPCR) (Fig. 1G). Notably, Notch2 deficiency also restored the anabolic response to mechanical overload and improved muscle hypertrophy in aged mice (Fig. S1A–C), suggesting that Notch2 is required for sarcopenia development and anabolic resistance in aged mice.

**Fig. 1.**
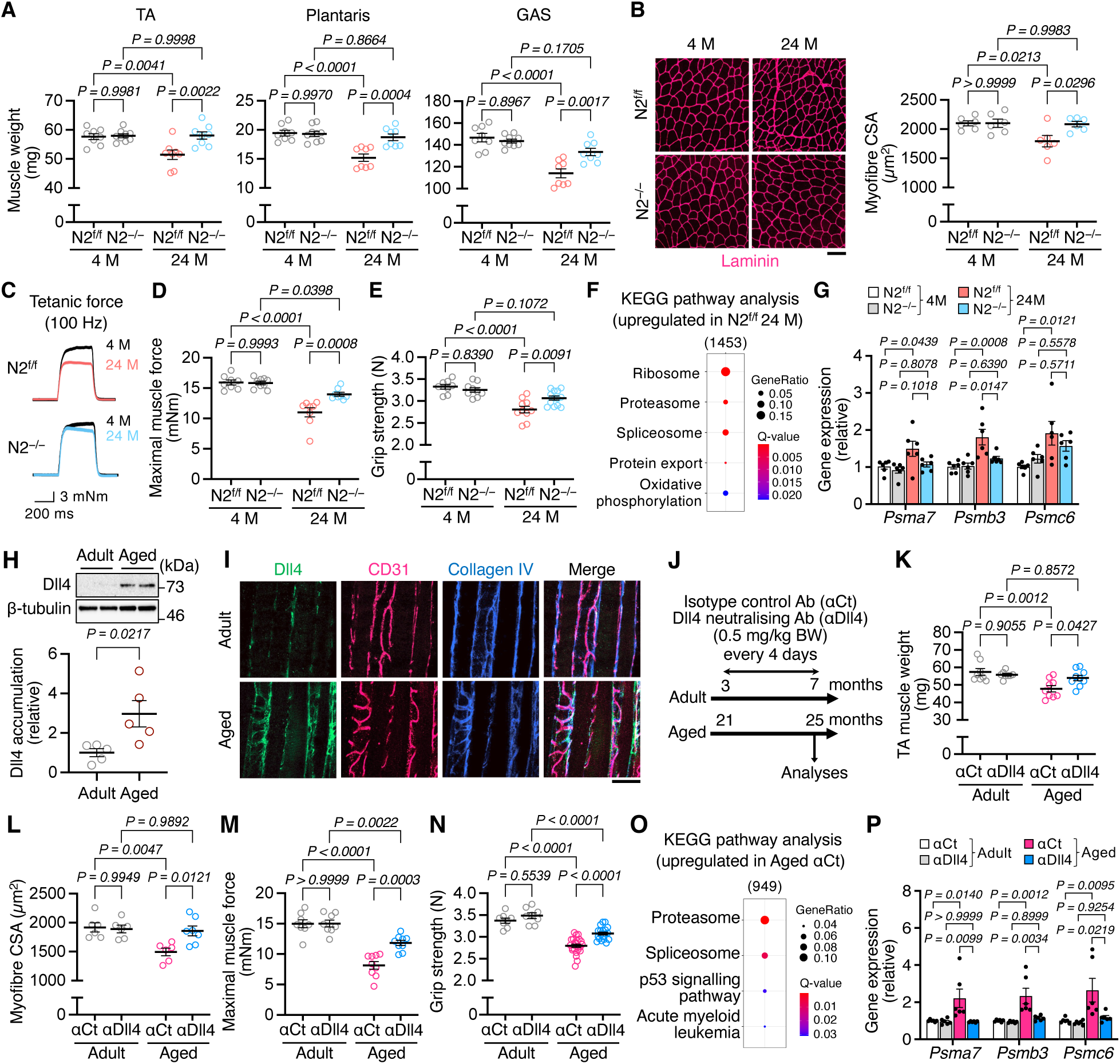
Blockade of the Dll4–Notch2 axis attenuates sarcopenic phenotypes. A, Muscle weights of tibialis anterior (TA), plantaris (PLA), and gastrocnemius (GAS) in *Notch2^flox/flox^* (N2^f/f^) and *Mlc1f^Cre/+^;Notch2^flox/flox^*(N2^−/−^) mice at 4 and 24 months of age (4 M and 24 M; n = 8). **B**, Representative images of TA muscle cross sections immunostained for laminin. Quantification of myofiber cross-sectional area (CSA; n = 6; scale bar, 100 µm). **C,D**, TA muscle force generation. Representative tetanic force traces and maximal muscle force (mNm; n = 8). **E**, Grip strength (n = 8– 13). **F**, Top five Kyoto Encyclopedia of Genes and Genomes (KEGG) pathways enriched in genes upregulated in 24-month N2^f/f^ mice in the RNA-seq (FC >1.2, q <0.05). **G,** qPCR analyses of proteasome-related genes (n = 6). **H**, Representative immunoblotting of Dll4 and β-tubulin (loading control) in plantaris muscles from adult (3–6 months) and aged (24–26 months) mice. Dll4 protein levels were quantified (n = 5). **I**, Representative images of TA muscle longitudinal sections immunostained for Dll4 (*green*), CD31 (*magenta*), and collagen IV (*blue*) (scale bar, 100 µm). **J–P**, Wild-type mice were administered anti-Dll4 neutralizing antibody (αDll4; 0.5 mg/kg) every 4 days for 4 months from 3 or 21 months of age; isotype control IgG (αCt) was used as a control. **j**, Experimental scheme. **K**, TA muscle weights (n = 8–9). **L**, Myofiber CSA (n = 6–7). **M**, TA maximal muscle force (n = 8–9). **N**, Grip strength (n = 8 adult, n = 19–24 aged). **O**, Top five KEGG pathways enriched in genes specifically upregulated in aged αCt mice in RNA-seq (FC >1.5, q <0.05). **P,** qPCR analyses of proteasome-related genes (n = 6). Data are shown as mean ±SEM. P values were determined by Student’s two-tailed unpaired t-test (**h**), and two-way ANOVA with Tukey’s post-hoc test (**A,B,D,E,G,K– N,P**).

Because Dll4 is predominantly expressed in vascular ECs in muscle (16) and serves as a ligand for myofiber Notch2 (19), we determined whether endothelial Dll4 contributes to muscle aging. Immunoblot analyses revealed markedly increased Dll4 protein levels in aged muscle (Fig. 1H), and Dll4 accumulation was mainly localized to the vascular and perivascular regions of aged muscle (Fig. 1I). To investigate the effects of attenuating Dll4 activity *in vivo*, we administered a Dll4-neutralizing antibody (αDll4) or an isotype control antibody (αCt) to young and aged mice at a dose of 0.5 mg/kg for four months (Fig. 1J). Systemic Dll4 attenuation substantially prevented age-associated declines in muscle mass, cross-sectional area (CSA), maximal force generation, and grip strength (Fig. 1K–N). Furthermore, αDll4 treatment improved both rotarod performance and treadmill endurance in aged mice, consistent with the findings observed in Notch2-deficient aged mice (Fig. S2A–D). These beneficial effects on muscle mass and function were also observed in aged mice treated with a lower dose of the Dll4-neutralizing antibody (0.1 mg/kg; Fig. S3A–E) and in geriatric mice in which treatment was initiated at 28 months of age (Fig. S3F–I). In contrast, αDll4 treatment did not affect muscle growth in young mice (0.1 and 0.5 mg/kg; Fig. S4A–C). Transcriptome analyses revealed age-dependent upregulation of proteasome-related genes, which were markedly suppressed by αDll4 (Fig. 1O,P), demonstrating endothelial Dll4 as a key regulator of muscle aging. These findings indicate that the endothelial Dll4–myofiber Notch2 axis plays a pivotal role in the development of sarcopenia. The beneficial effects of blocking either Dll4 or Notch2 on muscle mass and function were not associated with changes in the abundance of muscle satellite cells, myonuclear number, NMJ morphology, or fiber-type composition (Fig. S5A–J), suggesting that the Dll4-Notch2 axis influences muscle aging without affecting these age-related parameters.

### Soluble Dll4 preferentially binds to collagen I and Notch2

Given that systemic Dll4 blockade by a neutralizing antibody markedly attenuated sarcopenic phenotypes, we further investigated the effects of an Fc-tagged recombinant soluble Dll4 protein (rDll4-Fc), a competitive inhibitor of endogenous Dll4 (19, 24), in aged mice. During the 2-month treatment period (Fig. 2A), grip strength progressively decreased in vehicle-treated aged mice but was largely preserved in aged mice receiving intraperitoneal rDll4-Fc (Fig. 2B). Consistently, rDll4-Fc-treated mice also exhibited greater muscle force generation and increased muscle mass compared with control mice (Fig. 2C,D), indicating that rDll4-Fc as well as αDll4 exerts protective effects against age-associated muscle deterioration. Single-nucleus RNA-seq (snRNA-seq) demonstrated that *Notch2*, but not other Notch receptors, was detectable in myonuclei and upregulated in aged muscle regardless of fiber type (Fig. 2E, Fig. S6A,B). The expression of *Foxo1* and *Foxo3a*, regulators of muscle atrophy (25), also increased in aged myofibers, but was suppressed by rDll4-Fc (Fig. 2E).

**Fig. 2.**
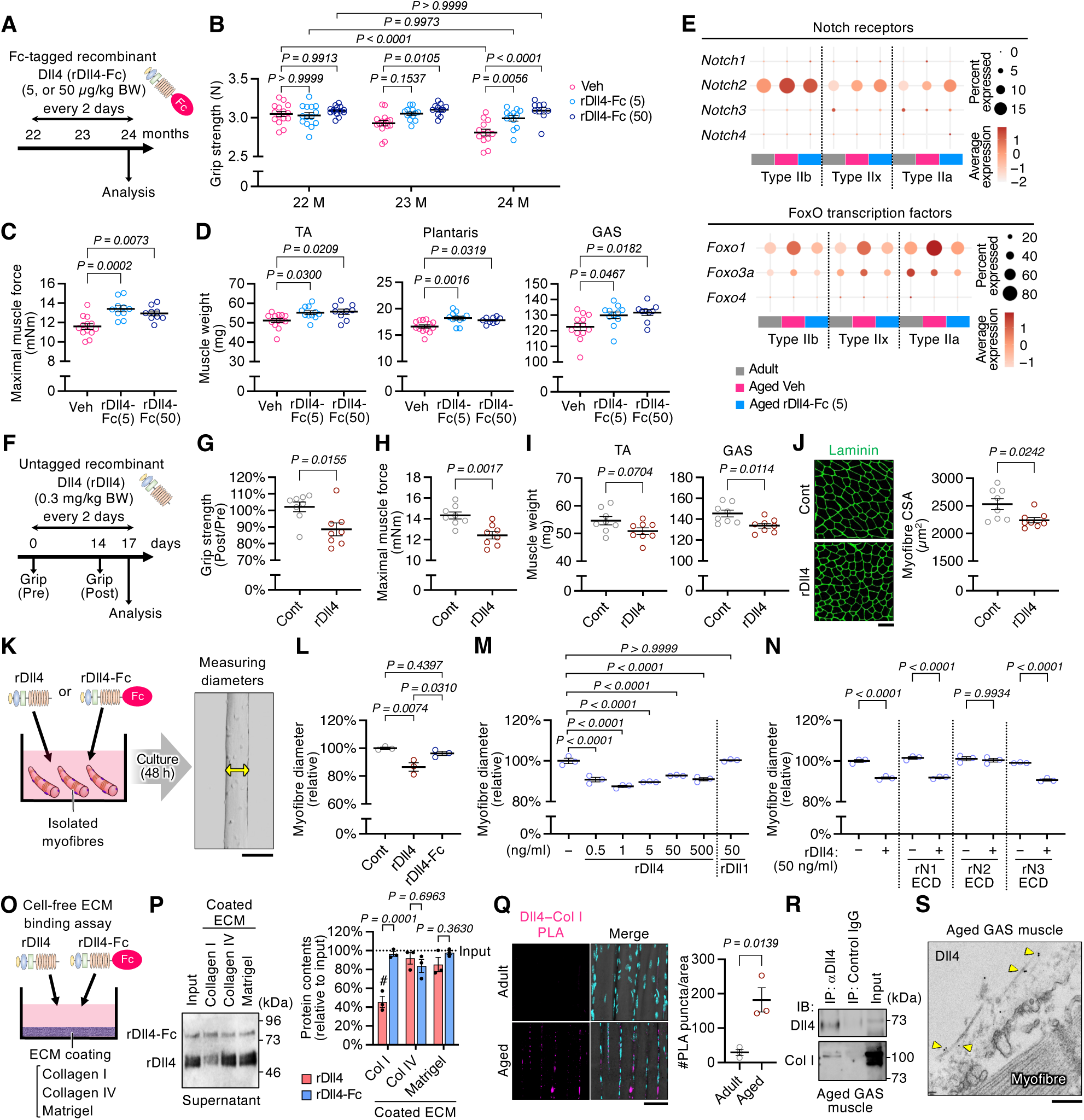
Soluble Dll4 binds to collagen I. A–E, Wild-type mice were injected with Fc-tagged recombinant Dll4 protein (rDll4-Fc; 5 or 50 µg/kg) every other day for 2 months from 22 months of age; vehicle (Veh) served as control. **A**, Experimental scheme. **B–D**, Grip strength (**B**), TA maximal muscle force (**C**), and muscle weights (**D**) (n = 9–15). **E**, snRNA-seq dot plots showing expression of Notch receptors and FoxO transcription factors in myonuclei. **F–J**, Wild-type mice (12 weeks old) were injected with untagged recombinant Dll4 protein (rDll4; 0.3 mg/kg) every other day for 17 days; vehicle was used as a control. **F**, Experimental scheme. **G–J**, Grip strength (**G**), TA maximal muscle force (**H**), muscle weights (**I**), and TA muscle cross-sections immunostained for laminin and myofiber CSA (**J**) (n = 8; scale bar, 100 µm). **K–N**, Effects of recombinant Dll4 on isolated myofibers in culture. **K**, Experimental scheme and representative images of myofibers cultured for 48 h (scale bar, 100 µm). **L–N**, Myofiber diameters after treatment with rDll4 or rDll4-Fc (50 ng/ml each) (**L**), rDll4 or rDll1 (**M**), or rDll4 (50 ng/ml) in combination with recombinant extracellular domain (ECD) of Notch1, Notch2, or Notch3 (500 ng/ml each) (**N**) (n = 3). **O, P**, Cell-free ECM-binding assay and quantification of unbound Dll4 proteins by immunoblotting (n = 4). **Q**, Representative TA longitudinal sections subjected to Dll4-collagen I proximity ligation assay (PLA), showing PLA puncta (*magenta*), DAPI (*cyan*), and corresponding transmitted light images, with PLA puncta quantification (scale bar, 50 µm). **R**, Representative immunoblot analysis of Dll4 and collagen I in anti-Dll4 immunoprecipitates from aged gastrocnemius (GAS) muscle lysates. **S**, Immunogold EM images of Dll4 in aged GAS. Yellow arrowheads indicate collagen-associated Dll4 (scale bar, 200 nm). Data are shown as mean ±SEM. P values were determined by two-way ANOVA with Tukey’s post-hoc test (**b**), one-way ANOVA with Tukey’s post-hoc test (**c,d,l–n**), Student’s two-tailed unpaired t-test (**G– J,Q**), and two-way ANOVA with Šídák’s multiple-comparisons test (**P**); #Significant difference from 100% input by one-sample t-test.

The activity of recombinant Dll4 is highly dependent on its physical form. While soluble rDll4-Fc acts as an antagonist (24), rDll4-Fc immobilized on culture dishes activates Notch receptors on plated cells, thereby recapitulating the activity of a membrane-bound Notch ligand (26). Unexpectedly, systemic administration of an untagged, soluble recombinant Dll4 protein (rDll4) to young mice induced muscle atrophy, accompanied by substantial reductions in grip strength, muscle force generation, muscle mass, and CSA (Fig. 2F–J). Consistently, treatment of isolated myofibers with soluble rDll4, but not soluble rDll4-Fc, markedly reduced myofiber size in culture (Fig. 2K,L). Soluble rDll4 induced myofiber atrophy over a broad concentration range (0.5–500 ng/ml), whereas an untagged, soluble recombinant Dll1 protein (rDll1) did not exert such an effect (Fig. 2M). Strikingly, rDll4-induced myofiber atrophy was completely abolished by the recombinant extracellular domain of Notch2 (rN2-ECD) fragment protein but not by the corresponding extracellular domains of Notch1 (rN1-ECD) or Notch3 (rN3-ECD) (Fig. 2N). These observations suggest that soluble rDll4 preferentially interacts with the extracellular domain of Notch2.

We previously found that Dll4 accumulates in the extracellular matrix of diabetic skeletal muscle (19), raising the possibility that extracellular matrix components provide a scaffold that facilitates soluble Dll4-mediated Notch activation. To test this, we examined the binding of soluble rDll4 and rDll4-Fc proteins to extracellular matrix components in a cell-free system (Fig. 2O). rDll4, but not rDll4-Fc, specifically bound to collagen I, whereas little or no binding was detected to collagen IV or Matrigel, which is composed primarily of laminin-111, collagen IV, and entactin (Fig. 2P). Remarkably, abundant Dll4–collagen I complexes were observed in aged muscle (Fig. 2Q,R, and Fig. S7A), and Dll4 protein localized along collagen-like fibers in the extracellular space (Fig. 2S). Together, these findings suggest that Dll4 binds to collagen I, which may act as a scaffold for soluble Dll4 and facilitate Notch2 activation in aged muscle.

### Soluble Dll4 drives endothelial senescence and vascular regression

Cellular senescence is associated with tissue aging and age-related functional decline. Having shown that Dll4 blockade reduced the expression of aging-related genes markedly and ameliorated sarcopenia (Fig. 1–2), we examined the effect of Dll4 inhibition on cellular senescence in muscle tissues. The abundance of SPiDER-βgal^+^ senescent cells increased with aging, but was substantially reduced by αDll4 treatment (Fig. 3A). snRNA-seq analysis further demonstrated that ECs, but not pericytes or fibro/adipogenic progenitors, exhibited the most pronounced age-associated increase in the senescence marker Cdkn1a (p21) (27, 28) (Fig. 3B). This induction was attenuated by soluble rDll4-Fc treatment (Fig. 3B). Co-immunostaining for the EC marker CD31 and SPiDER-βgal showed that the number of senescent ECs decreased following soluble rDll4-Fc treatment compared with that in vehicle controls of aged muscle (Fig. 3c,D), while sustained exposure to rDll4 in young mice was sufficient to increase the number of senescent cells (Fig. 3D). We next evaluated whether soluble rDll4 directly induces endothelial senescence and found that prolonged exposure of human umbilical vein endothelial cells (HUVECs) to soluble rDll4 increased the proportion of SPiDER-β-gal^+^ cells, accompanied by upregulation of senescence-associated genes, whereas co-treatment with either a Dll4-neutralizing antibody or soluble rDll4-Fc largely prevented these responses (Fig. 3E–G).

**Fig. 3.**
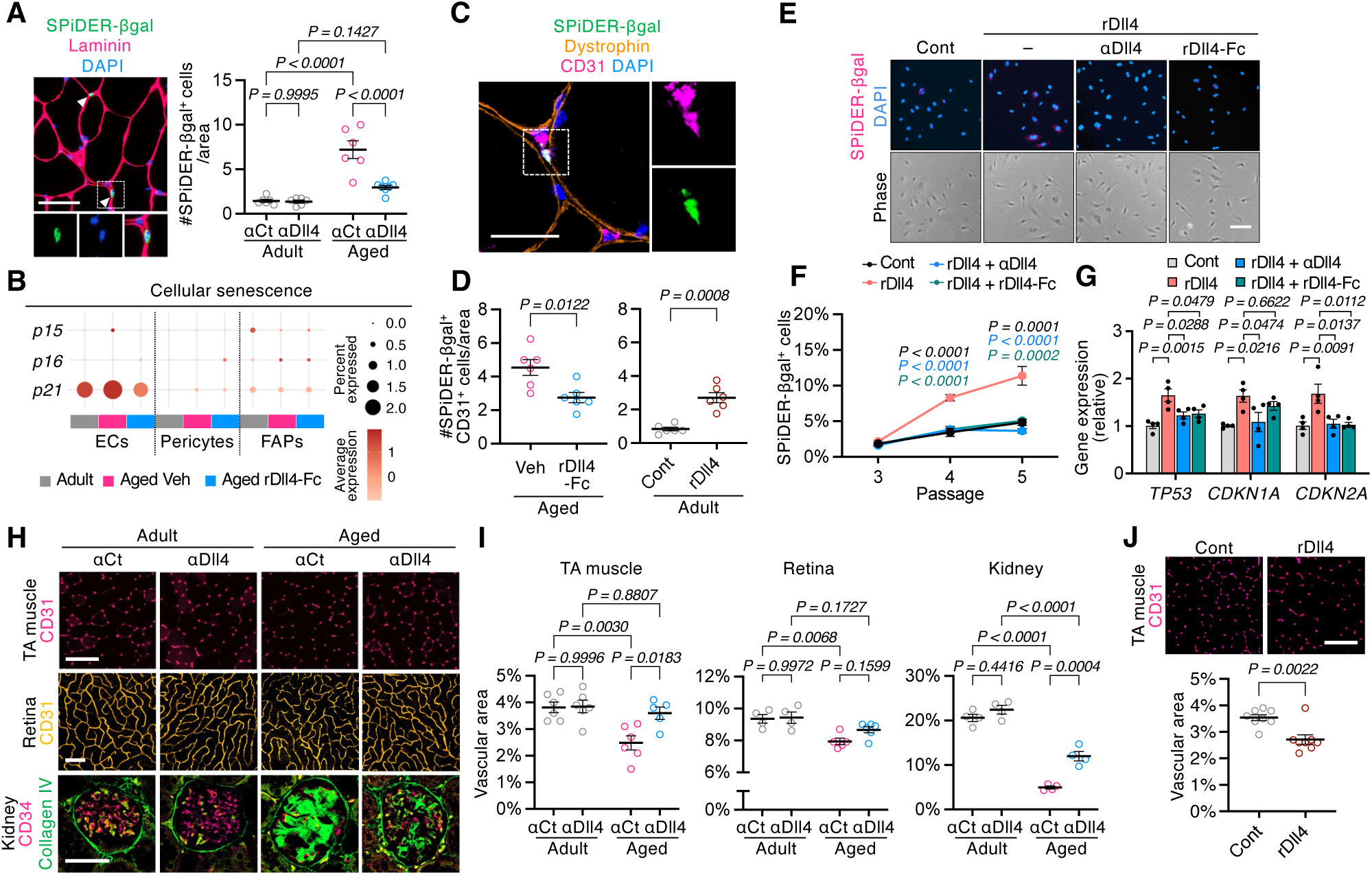
Soluble Dll4 induces endothelial senescence. A, Representative images of tibialis anterior (TA) muscle cross sections stained with SPiDER-β-gal (*green*), laminin (*magenta*), and DAPI (*blue*), with quantification of senescent cells (n = 6–7; scale bar, 50 µm). **B**, snRNA-seq dot plots of senescence-related genes in endothelial cells (ECs), pericytes, and fibro-adipogenic progenitors (FAPs) from adult and aged mice treated with rDll4-Fc or vehicle. **C**, Representative images of TA muscle cross sections stained for SPiDER-βgal (*green*), CD31 (*magenta*), dystrophin (*orange*), and DAPI (scale bar, 20 µm). **D**, Number of senescent ECs in aged mice (24 months old) treated with vehicle or rDll4-Fc (5 µg/kg) and adult mice (12 weeks old) treated with vehicle or rDll4 (0.3 mg/kg) (n = 6). **E**, Representative SPiDER-βgal staining with DAPI and phase-contrast images of human umbilical vein endothelial cells (HUVECs) cultured under the indicated conditions for 9 days across passages 3–5 (scale bar, 100 µm). **F**, Quantification of SPiDER-βgal positivity at each passage (n = 4). **G**, qPCR analysis of senescence-related genes in HUVECs (n = 4). **H**, Representative TA and retina sections stained for CD31 (*magenta* and *yellow*, respectively), and kidney sections stained for CD34 (*magenta*) and collagen IV (*green*), from adult and aged mice treated with αDll4 or αCt (scale bar, 100 µm for TA and retina; 50 µm for kidney). **I**, Vascular area in each tissue (n = 4–6). **J**, Representative TA cross sections from rDll4-treated mice stained for CD31 (*magenta*), with quantification of vascular area (n = 8; scale bar, 100 µm). Data are shown as mean ±SEM. P values were determined by Student’s two-tailed unpaired t-test (**D,J**), one-way ANOVA with Tukey’s post-hoc test (**F,G**), and two-way ANOVA with Tukey’s post-hoc test (**A,I**).

Because endothelial senescence is closely linked to capillary rarefaction during aging, we next examined the systemic effect of Dll4 blockade on vasculature in multiple tissues. Capillary densities in skeletal muscle, retina, and kidney were substantially reduced in aged mice compared with those in young mice (Fig. 3H,I). Remarkably, chronic administration of αDll4 from 21 months of age largely prevented this age-associated vascular regression (Fig. 3H,I). Conversely, systemic administration of soluble rDll4 to young mice was sufficient to reduce vascular density (Fig. 3J). Together, these findings demonstrate soluble Dll4 as a direct inducer of endothelial senescence and vascular regression in aging.

### Systemic Dll4 inhibition promotes healthy longevity

Based on the finding that Dll4 blockade prevents vascular regression in multiple tissues (Fig. 3H-I), we next examined whether reducing Dll4 activity confers broader benefits on systemic aging (Fig. 4A). Aging is associated with a decline in tissue NAD^+^ levels, including in the liver (29). Notably, administration of a Dll4-neutralizing antibody markedly restored hepatic NAD^+^ levels in aged mice and reduced the number of SPiDER-βgal^+^ senescent cells in the liver (Fig. 4B,C). Histological analyses demonstrated that Dll4 inhibition ameliorated age-associated renal pathological changes, including Bowman’s capsule enlargement, tubular injury, and interstitial fibrosis (Fig. 4D–G, Fig. S8A). Bulk RNA-seq to further characterize the systemic effects of Dll4 blockade revealed that aging upregulated genes involved in arachidonic acid metabolism in the liver and retinol metabolism in the kidney, whereas Dll4 blockade suppressed these changes (Fig. 4 H–M). We also identified three genes, *Chil3*, *Gm27252*, and *Cirbp*, that were commonly upregulated during aging and downregulated by Dll4 blockade across muscle, liver, and kidney (Fig. 4N). qPCR analysis confirmed the age-associated induction of *Cirbp* and its suppression by αDll4 treatment (Fig. 4O), suggesting that *Cirbp* is a potential molecular signature of systemic tissue aging.

**Fig. 4.**
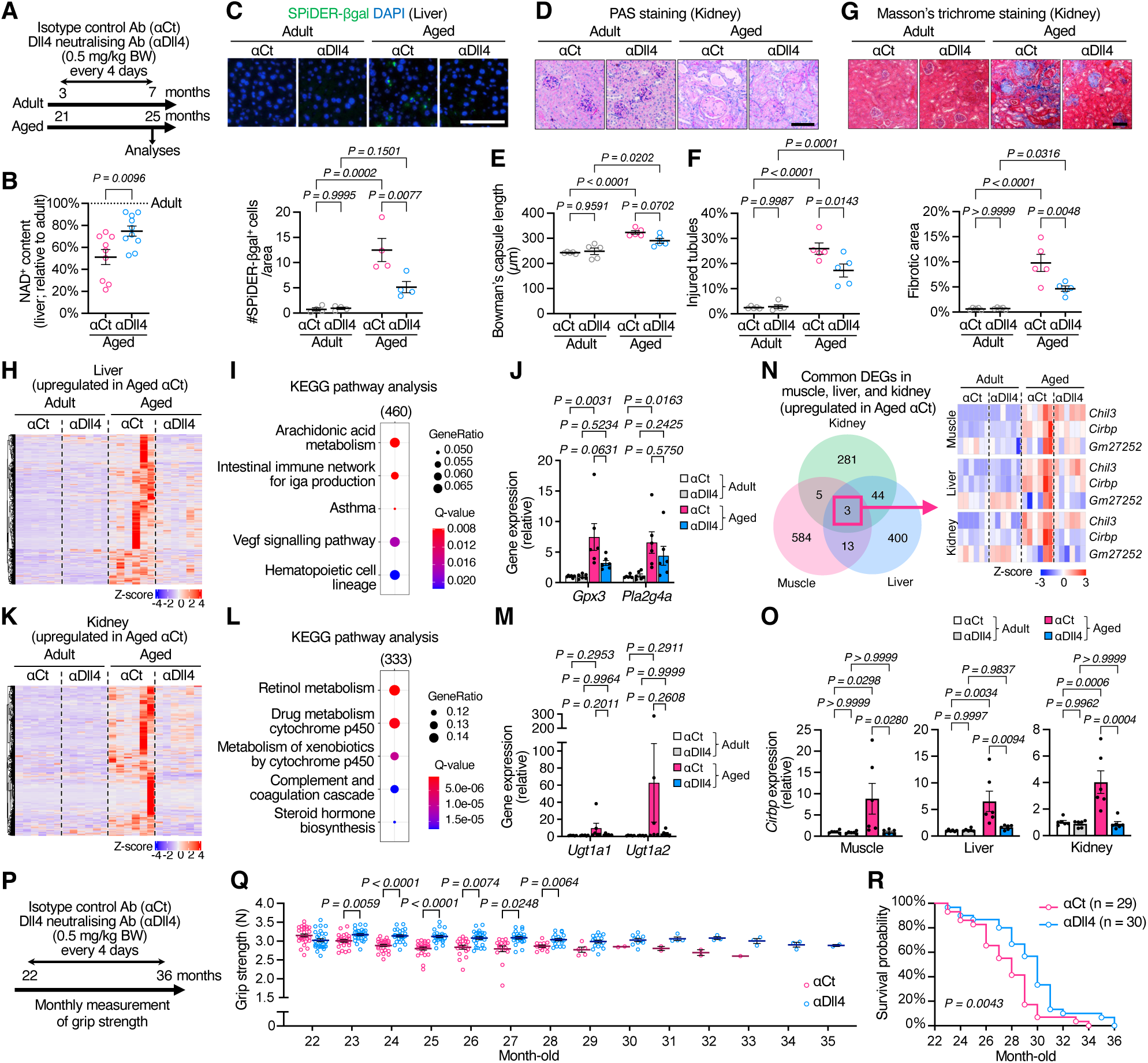
Systemic inhibition of Dll4 extends healthspan and lifespan. A–O, Wild-type mice were administered αDll4 (0.5 mg/kg) every 4 days for 4 months from 3 or 21 months of age; αCt served as a control. **A**, Experimental scheme. **B**, Liver NAD^+^ content (n = 9–10). **C**, Representative liver sections stained for SPiDER-βgal (*green*) and DAPI, with quantification of senescent cells (n = 4; scale bar, 100 µm). **D–F**, Representative PAS-stained kidney sections (**D**), with quantification of Bowman’s capsule length (**E**) and the percentage of injured tubules (**F**) (n = 5; scale bar, 100 µm). **G**, Representative Masson’s trichrome-stained kidney sections, with quantification of fibrotic area (n = 5; scale bar, 100 µm). **H,I**, Heatmap and top five Kyoto Encyclopedia of Genes and Genomes (KEGG) pathways enriched in genes specifically upregulated in liver from aged αCt mice (FC >1.5, q <0.05) in RNA-seq. **J**, qPCR analyses of arachidonic acid metabolism-related genes (n = 6). **K,L,** Heatmap and top five KEGG pathways enriched in genes specifically upregulated in kidney from aged αCt mice (FC >1.5, q <0.05) in RNA-seq. **M**, qPCR analyses of genes shared across the top enriched KEGG pathways (n = 6). **N**, Venn diagram of genes upregulated in muscle, liver, and kidney of aged αCt mice, and heatmap of three genes commonly upregulated across the tissues. **O**, qPCR analyses of *Cirbp* in each tissue (n=6). **P–R**, Long-term effects of Dll4 inhibition on aged mice. **P**, Experimental scheme. **Q**, Monthly grip strength monitoring (initial n = 29–30/group). **R**, Survival curve. Data are shown as mean ±SEM. P values were determined by Student’s two-tailed unpaired t-test (**B**), two-way ANOVA with Tukey’s post-hoc test (**C,E–G,J,M,O**), and mixed-effects model (REML) with Šídák’s post-hoc test (**Q**). P value of the survival experiment (**R**) was determined by the log-rank test.

We next examined whether the effects of chronic Dll4 inhibition can translate into prolonged healthy lifespan. Grip strength is a non-invasive surrogate measure of systemic muscle function and declines in parallel with other measures of physical function in mice (30). Administration of αDll4 was initiated in mice at 22 months of age, and grip strength was monitored longitudinally until the end of life (Fig. 4P). Strikingly, αDll4-treated mice maintained grip strength levels throughout life compared with αCt-treated mice, which exhibited the expected progressive functional decline (Fig. 4Q). Kaplan– Meier analysis further demonstrated that systemic Dll4 inhibition extended survival in aged mice relative to that in control-treated mice (Fig. 4R). Hence, Dll4 acts as a systemic regulator of biological aging, and attenuation of excessive Dll4 signaling promotes healthy aging and extends both healthspan and lifespan.

### Endothelial cells release full-length Dll4 via a migration-associated mechanism

Dll4 has been reported to be released into the extracellular space and detected in the circulation (19, 31–34). We therefore examined circulating Dll4 levels during aging and found that serum Dll4 concentrations were markedly higher in aged mice than in young controls (Fig. 5A). We next sought to identify the stimuli that promote Dll4 release from ECs. snRNA-seq revealed an enrichment of inflammatory cytokine response pathways in ECs from aged muscle (Fig. 5B). Inflammatory stimuli, such as lipopolysaccharide (LPS) can induce cellular senescence and upregulate Dll4 expression in ECs (35, 36). Consistent with a potential role for inflammation in Dll4 release, systemic LPS administration increased circulating Dll4 levels in adult mice (Fig. 5C). In addition to inflammatory stimuli, hypoxia upregulates Dll4 expression and induces cellular senescence in ECs (37, 38). To identify stimuli that directly induce Dll4 release from ECs, we treated HUVECs with LPS, tumor necrosis factor-α, or interleukin-1β as inflammatory stimuli, or with cobalt chloride to mimic hypoxic conditions. Immunoblot analysis revealed increased Dll4 levels in extracellular vesicle (EV)-depleted conditioned media following exposure to each of these stimuli (Fig. 5D), suggesting that inflammatory and hypoxia-mimetic stimuli promote Dll4 release from ECs.

**Fig. 5.**
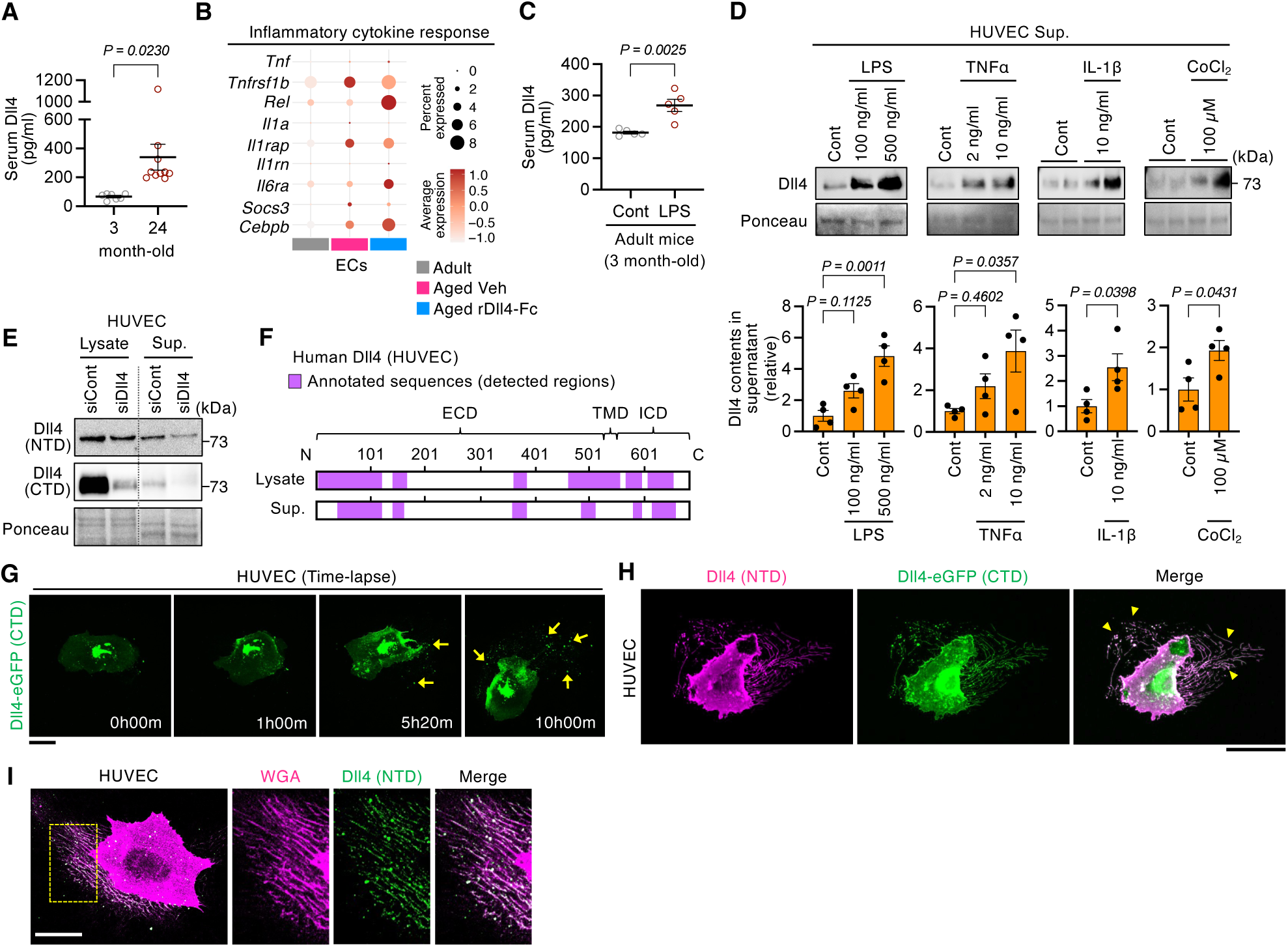
Full-length Dll4 is released from endothelial cells in response to inflammatory stimuli. A, Serum Dll4 levels in 3- and 24-month-old mice (n = 7–10). **B**, snRNA-seq dot plots showing the expression of inflammatory cytokine response-related genes in ECs from adult and aged mice treated with rDll4-Fc or vehicle for 2 months. **C**, Serum Dll4 levels 3 h after lipopolysaccharide (LPS; 1 mg/kg) administration in 3-month-old mice (n = 5). **D**, Immunoblotting and quantification of Dll4 in extracellular vesicle (EV)-depleted supernatants from HUVECs treated with LPS (100 or 500 ng/ml), tumor necrosis factor (TNF)-α (2 or 10 ng/ml), interleukin (IL)-1β (10 ng/ml), or cobalt chloride (CoCl_2_, 100 µM) for 24 h (n = 4). Ponceau staining was used as a loading control. **E**, Immunoblotting of Dll4 in HUVEC lysates and EV-depleted supernatants following transfection with Dll4-targeting siRNA (siDll4) or control siRNA (siCont), using antibodies against the N-or C-terminus of Dll4. **F**, Dll4 protein regions identified by liquid chromatography–tandem mass spectrometry (LC–MS/MS) in anti-Dll4 immunoprecipitates from HUVEC lysates and supernatants. **G**, Time-lapse fluorescence images of HUVECs expressing C-terminally eGFP-tagged Dll4 (Dll4–eGFP (CTD); *green*) over a 10-h period. Yellow arrows indicate released Dll4 (scale bar, 50 µm). **H**, Fluorescence images of HUVECs expressing Dll4–eGFP (CTD; *green*) and immunostained for Dll4 N-terminus (NTD; *magenta*). Yellow arrowheads indicate retraction fibers and migrasome-like vesicles (scale bar, 50 µm). **I**, Representative images of HUVECs stained for endogenous Dll4 (NTD; *green*) and WGA (*magenta*), with corresponding magnified views (scale bar, 50 µm). Yellow arrowheads indicate retraction fibers and migrasome-like vesicles (scale bar, 50 µm). Data are shown as mean ± SEM. P values were determined by Student’s two-tailed unpaired t-test (**A,C,D** [two right panels]), and one-way ANOVA with Tukey’s post-hoc test (**D** [two left panels]).

We further characterized the molecular form of extracellular Dll4. Immunoblot analysis using antibodies against either the N-or C-terminal domain of Dll4 detected bands of comparable molecular weight in EV-depleted conditioned media and cell lysates from cultured HUVECs (Fig. 5E). These bands were diminished following siRNA-mediated Dll4 knockdown, confirming their specificity (Fig. 5E). Furthermore, liquid chromatography–tandem mass spectrometry (LC–MS/MS) analysis of EV-depleted conditioned media identified Dll4-derived peptides spanning multiple regions of the protein, including its N- and C-terminal regions (Fig. 5F). Collectively, these findings support the presence of full-length Dll4 in the extracellular milieu.

To visualize Dll4 protein dynamics, we performed time-lapse live-cell imaging of HUVECs expressing Dll4 fused to green fluorescent protein (GFP) at its intracellular C-terminus. Notably, Dll4 accumulated in membrane-associated fragments and particle-like structures that were deposited and released during EC migration (Fig. 5G). Immunofluorescence staining with an antibody recognizing the N-terminal domain of Dll4, together with detection of fluorescence from the C-terminal GFP tag, demonstrated the presence of both regions within these extracellular structures (Fig. 5H). These observations further support the release of Dll4 in a form retaining both its N- and C-terminal regions. We next characterized these Dll4-containing structures. Their morphology resembled that of migrasomes (39), migration-associated organelles formed on retraction fibers during cell movement. Consistent with this possibility, Dll4 colocalized with wheat germ agglutinin (WGA) staining, which is used to visualize migrasomes and retraction fibers (40) (Fig. 5I). As migrasomes can release their contents into the extracellular environment (39), these observations suggest that migrasome-associated structures, including retraction fibers, may participate in the release of full-length Dll4 from ECs. This mechanism may contribute, at least in part, to the extracellular accumulation of Dll4 during aging.

### Kidney-derived circulating Dll4 induces muscle wasting

Having shown that serum Dll4 levels were elevated in aged mice (Fig. 5A), we next investigated the relationship between circulating Dll4 levels and pathophysiological states in humans. Serum Dll4 concentrations were measured in a cohort of 1,449 individuals aged 65 years or older and compared with biochemical and physiological parameters. Circulating Dll4 exhibited only a weak correlation with chronological age (ρ = 0.2385) (Fig. 6A and Table S1), suggesting that factors other than age per se may determine Dll4 abundance in elderly individuals. Among all clinical parameters examined, estimated glomerular filtration rate (eGFR) showed the strongest association with circulating Dll4 levels (ρ = −0.3952) (Fig. 6B and Tables S1–3), implicating renal dysfunction as a major determinant of systemic Dll4 elevation.

**Fig. 6.**
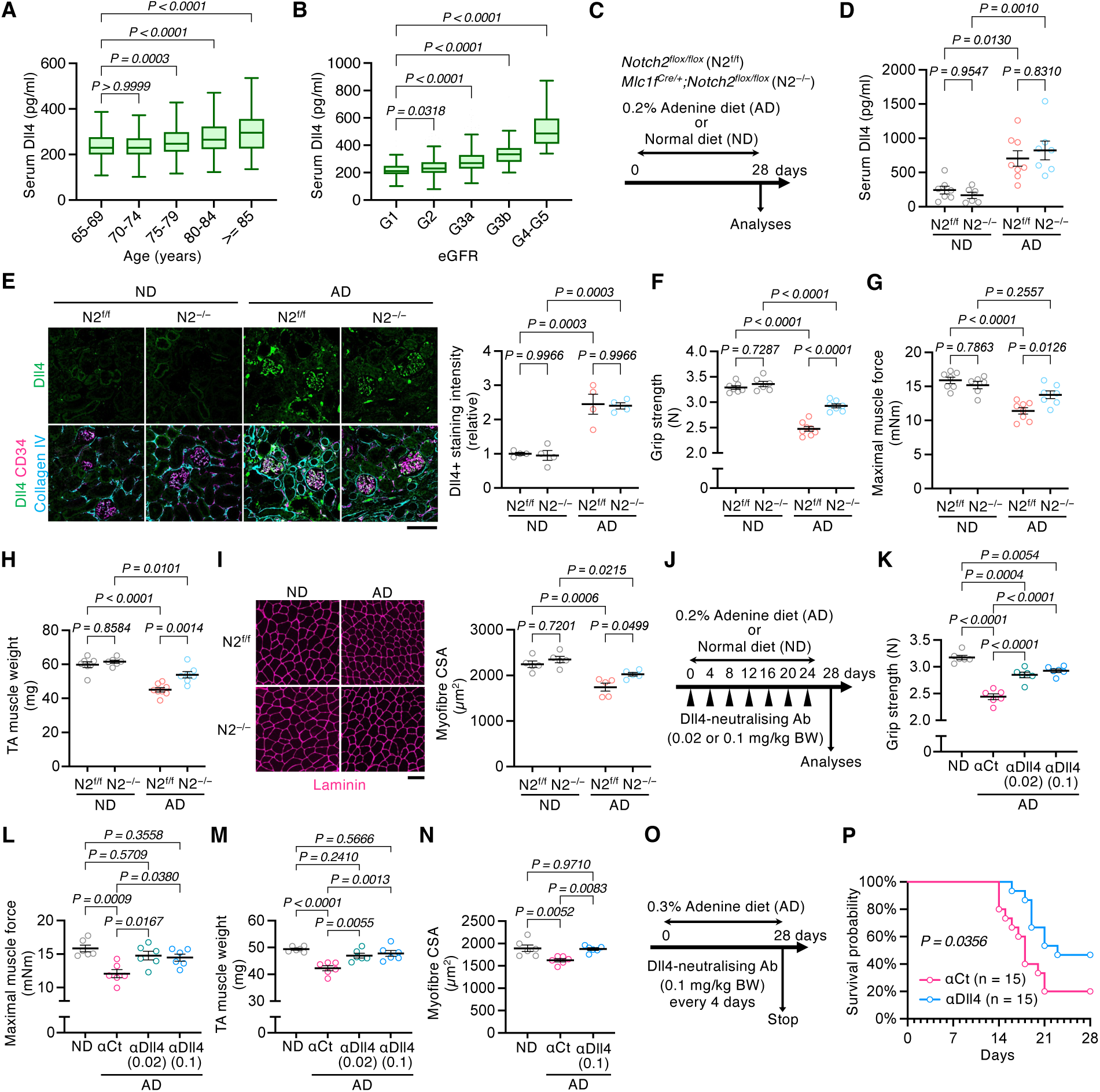
Circulating Dll4 mediates muscle atrophy in chronic kidney dysfunction. A,B, Serum Dll4 concentrations according to age (**A**) and kidney function (eGFR) (**B**) in human subjects. **C–I**, N2^f/f^ and N2^−/−^ mice (8–12 weeks old) were fed a 0.2 % adenine diet (AD) for 28 days. **C**, Experimental scheme. **D**, Serum Dll4 levels (n = 6–8). **E**, Kidney tissue sections stained for Dll4 (*green*), CD34 (*magenta*), and collagen IV (*cyan*), with quantification of Dll4^+^ fluorescence intensity. **F–I**, Grip strength (**F**), muscle force (**G**), muscle weights (**H**), and tibialis anterior (TA) cross sections stained for laminin, with myofiber cross-sectional area (CSA) quantification (**I**) (n = 6–8; scale bar, 100 µm). (**J–N**, Wild-type mice (12 weeks old) were injected with αDll4 (0.02 or 0.1 mg/kg) every 4 days during AD feeding. **J**, Experimental scheme. **K–N**, Grip strength (**K**), muscle force (**L**), muscle weights (**M**), and TA cross sections stained for laminin, with myofiber CSA quantification (**N**) (n = 6). **O,P**, Experimental scheme and survival curve. P values were determined by two-way ANOVA with Tukey’s post hoc test (**D–I**), and one-way ANOVA with Tukey’s post-hoc test (**K–N**). P value of the survival experiment (**P**) was determined by the log-rank test. Data are shown as mean ± SEM unless otherwise indicated. Human cohort data (**A,B**) are shown as Tukey box plots and were analyzed using the Kruskal–Wallis test.

Chronic kidney disease (CKD) is associated with muscle weakness, leading to secondary sarcopenia (41). We further examined whether renal dysfunction promotes sarcopenia through the Dll4–Notch2 axis. To this end, we used an adenine diet-induced CKD model and assessed circulating Dll4 levels and muscle phenotypes in mice with myofiber-specific deletion of Notch2 (Fig. 6C). Serum Dll4 concentrations were remarkably elevated in CKD mice irrespective of genotypes (Fig. 6D), concomitant with increased expression of renal injury markers (Fig. S9A–C). This increase was unlikely to result from CKD-induced impairment of renal filtration, as urinary Dll4 levels remained unchanged in CKD mice, a finding that was also observed in aged mice (Fig. S10A,B). Immunohistochemical analyses further revealed robust Dll4 accumulation in glomerular and tubular ECs in CKD kidneys (Fig. 6E), suggesting that the injured kidney may serve as a major source of elevated circulating Dll4. Despite comparable degrees of renal dysfunction and similar serum Dll4 levels, mice with myofiber-specific Notch2 deletion were substantially protected from CKD-associated muscle deterioration, exhibiting greater muscle mass, force generation, and grip strength than wild-type mice with CKD (Fig. 6F–I). Moreover, systemic administration of a Dll4-neutralizing antibody markedly attenuated CKD-induced reductions in muscle mass and strength without affecting the underlying renal pathology (Fig. 6J–N and Fig. S9D–F). We assessed whether Dll4 blockade confers broader systemic benefits in severe CKD. Strikingly, Dll4 neutralization prolonged survival in mice with advanced CKD (Fig. 6O,P). Altogether, these findings suggest that circulating Dll4 acts as an endothelium-derived systemic factor that links renal dysfunction to muscle wasting through a kidney– vascular–muscle axis.

## Discussion

Here, we demonstrated that inhibition of the Dll4–Notch2 axis markedly attenuated age-related declines in muscle mass and function, despite failing to reverse several hallmarks of muscle aging, including reductions in satellite cell abundance, myonuclear number, and myofiber number, increased NMJ fragmentation, and a shift from fast to slow MyHC isoforms (3, 4). These findings suggest that the endothelial Dll4–myofiber Notch2 axis contributes substantially to sarcopenia development through mechanisms that are, at least in part, distinct from those underlying other established features of muscle aging.

Vascular endothelial dysfunction is a major determinant of age-related morbidity and mortality (1, 42). Accumulating evidence suggests that capillary aging is not merely a consequence of tissue aging but may also serve as an upstream driver of systemic aging (43, 44). In particular, age-associated endothelial senescence impairs angiogenic capacity and promotes progressive capillary rarefaction, resulting in reduced tissue perfusion, chronic inflammation, and organ dysfunction (28, 42). Sustained VEGF-A signaling counteracted age-associated capillary rarefaction and endothelial senescence, ameliorated multiple systemic manifestations of aging, and extended lifespan in mice (44). Together, these findings highlight the critical role of vascular integrity in maintaining tissue homeostasis and suggest that preventing microvascular deterioration may represent a promising strategy for extending healthspan. Herein, we demonstrated that prolonged inhibition of Dll4 activity markedly attenuated vascular regression, ameliorated age-related tissue alterations, and extended both healthspan and lifespan in aged mice. Conversely, systemic administration of soluble rDll4 induced endothelial senescence and vascular regression across multiple organs, even in young mice. Dll4 plays an essential role in vascular development during embryogenesis (45, 46) and in regulating vascular formation and remodeling in postnatal and adult tissues (33, 47–49). Indeed, Dll4 blockade has been extensively investigated as an anti-angiogenic cancer therapy because it induces excessive but nonproductive angiogenesis, thereby impairing functional tumor vascularization (50, 51). Despite the established importance of Dll4 signaling in vascular homeostasis and the potential adverse effects associated with its inhibition, we observed no overt adverse effects in aged mice treated with a Dll4-blocking antibody. Instead, Dll4 blockade ameliorated age-associated vascular abnormalities. One possible explanation for this apparent discrepancy is the difference in the extent of Dll4 inhibition. Here, the antibody was administered at relatively low doses (0.1–0.5 mg/kg) compared with the higher doses (approximately 5 mg/kg) used in previous studies (50, 51). We therefore speculate that partial inhibition of aberrantly elevated Dll4 signaling during aging may be sufficient to ameliorate age-associated vascular dysfunction while minimizing the adverse effects of excessive Dll4 blockade, consistent with the dose-dependent effects of Dll4 inhibition observed in wound healing (52). These findings suggest that controlled inhibition of aberrantly elevated Dll4 signaling may represent a promising therapeutic strategy for preserving microvascular integrity during aging.

Our functional comparison of soluble rDll4 proteins with and without an Fc tag revealed that collagen I acts as an extracellular scaffold that enables soluble rDll4 to selectively activate Notch2, but not Notch1 or Notch3. Notably, soluble rDll1 failed to induce myofiber atrophy despite the structural similarities between Dll1 and Dll4 (7, 8), suggesting that this activity is specific to Dll4. Canonical Notch activation generally requires membrane-bound ligands, as ligand endocytosis generates the mechanical pulling force necessary for receptor activation (7, 8). However, previous studies have reported that Jagged1 can be released in a soluble form through ADAM17-mediated proteolytic cleavage and subsequently activate Notch1 signaling (53, 54). Furthermore, a recent study using DNA origami-based nanopatterning demonstrated that prolonged interactions between soluble Jagged1 and Notch receptors can promote Notch activation in the absence of conventional ligand-mediated pulling forces or direct cell–cell contact (55). Together, our findings support a model in which collagen I serves as an extracellular scaffold that immobilizes soluble Dll4 and increases its local availability, thereby facilitating Notch2 activation through a mechanism distinct from canonical ligand endocytosis. Such extracellular immobilization may functionally mimic membrane-bound Notch ligands without requiring conventional ligand endocytosis, providing a potential explanation for the context-dependent effects of soluble Dll4 on vascular remodeling, endothelial senescence, and muscle aging. We also found that Dll4 is released in its full-length form rather than through proteolytic cleavage. Our immunocytochemical and time-lapse analyses revealed that full-length Dll4 localizes to migrasome-associated structures (39, 40), including retraction fibers, which extend during EC migration, subsequently collapse, and ultimately release Dll4-containing material into the extracellular space. Given that migrasomes can release their contents into the extracellular environment (39, 40), these observations suggest that migrasome-associated structures may contribute to the release of full-length Dll4. Collectively, our findings describe a previously unrecognized mode of endothelial Dll4 release associated with migrasome-like structures formed during EC migration. Further studies are needed to determine the precise mechanisms underlying this process and whether additional pathways contribute to Dll4 release.

CKD is associated with increased morbidity and mortality and accelerated tissue aging, driven by multiple CKD-related abnormalities, including metabolic acidosis, chronic inflammation, insulin resistance, elevated glucocorticoid levels, and renin–angiotensin system activation (56). Moreover, CKD is frequently accompanied by progressive muscle wasting, contributing to secondary sarcopenia development (41). Accordingly, CKD-associated muscle wasting is thought to arise from the interplay of multiple metabolic and inflammatory abnormalities rather than from a single pathogenic pathway (41, 56). Here, we found that circulating Dll4 levels were associated with renal dysfunction in both human and mouse CKD cohorts and that inhibition of the Dll4–Notch2 axis substantially ameliorated CKD-associated declines in muscle mass and strength. Furthermore, Dll4 inhibition improved survival in CKD mice. These findings suggest that targeting Dll4 may confer systemic benefits beyond the preservation of muscle mass and function in CKD.

Although aging manifests differently across organs, alterations in the systemic circulatory milieu have emerged as important determinants of biological aging. Heterochronic parabiosis studies have demonstrated that exposure to a young circulatory environment can restore regenerative capacity in aged tissues (57), whereas exposure to aged blood can rapidly induce cellular senescence and impair tissue regeneration (58, 59). These findings highlight the importance of circulating factors in the systemic regulation of aging, although the molecular mediators responsible for these effects remain incompletely defined. In this context, our findings suggest that circulating Dll4, whose levels increase with age, may represent a mediator of systemic aging.

Our multi-tissue transcriptomic analysis further identified *Cirbp* as a potential shared-tissue biomarker of aging-associated changes, as its expression increased with age and was suppressed by Dll4 inhibition in skeletal muscle, liver, and kidney. Thus, *Cirbp* may serve as a pharmacodynamic marker of systemic tissue stress and responses to Dll4-targeted interventions. As CIRBP is a stress-responsive RNA-binding protein (60), its induction may reflect a shared cellular response to hypoxia, inflammation, or metabolic stress and could potentially contribute to aging-associated phenotypes (61).

In summary, we identified soluble, full-length Dll4 as a potential common mediator of both primary and secondary sarcopenia. Notably, Dll4 blockade improved muscle function even when initiated in old or geriatric mice and extended both functional healthspan and lifespan. These findings highlight the Dll4–Notch2 axis as a promising therapeutic target for preserving vascular integrity, maintaining inter-organ homeostasis, and promoting healthy aging.

## Methods

### Animals

All animal experiments were approved by the Institutional Animal Care and Usage Committees of Kumamoto University (A2022-075, A2024-096) and the Tokyo Metropolitan Institute for Geriatrics and Gerontology (25008). The animals were housed in a specific pathogen-free environment at 22±2 °C and 40–80% humidity on a 12-h light cycle, with free access to food and water. Wild-type C57BL/6J mice were obtained from the Jackson Laboratory, Japan or TMIG. Wild-type C57BL/6N mice used in Figs. 1I, 2Q, 5A, S5A, and S10A were obtained from TMIG, Japan. *Notch2*-floxed mice (stock no. 010525) (20) were obtained from the Jackson Laboratory. *Mlc1f^Cre^* mice were kindly provided by S. Burden (21). *Mlc1f^Cre^* mice were crossed with *Notch2*-floxed mice to generate *Mlc1f^Cre/+^; Notch2^flox/flox^* (termed N2^−/−^) and control Cre-negative littermates (termed N2^f/f^) as described previously (19). Male mice were used in all experiments, and the ages of mice are described in each figure legend. For the systemic inflammation model, mice were intraperitoneally injected with lipopolysaccharide (LPS) (1 mg/kg), and blood was collected 3 h later for serum Dll4 measurement by enzyme-linked immunosorbent assay (ELISA).

### Neutralizing antibody and recombinant protein treatments *in vivo*

To examine the effect of inhibition of the Dll4 signaling pathway *in vivo*, wild-type mice were used. Ultra-LEAF Armenian hamster anti-Dll4 antibody (Dll4 Neutralizing antibody, αDll4; 130802, clone HMD4-1, BioLegend) or Ultra-LEAF Armenian hamster IgG control antibody (control antibody, αCt; 40968, clone HTK8881, BioLegend) was intraperitoneally administered every 4 days in each experiment. The dosage is described in each figure legend. Recombinant Dll4-Fc chimera protein (rDll4-Fc; 10089-D4, R&D Systems) was dissolved in phosphate-buffered saline (PBS) and administered (5 or 50 µg/kg body weight) every 2 days for 2 months. In contrast, to activate this signaling, untagged recombinant Dll4 protein (rDll4; 140-07, Thermo Fisher Scientific) was dissolved in PBS and administered (0.3 mg/kg body weight) every 2 days for 17 days. Control mice were intraperitoneally injected with an equivalent volume of vehicle.

### Chronic kidney disease (CKD) mouse model

After a 3-day acclimation to a powdered diet, mice were fed a 0.2% adenine diet (AD) or normal powdered diet for 4 weeks. For the survival experiment, 30 wild-type mice were divided into two groups, and starting at the initiation of a 0.3% AD, were intraperitoneally injected every 4 days with either αDll4 (0.1 mg/kg) or αCt (0.1 mg/kg). CKD mice were monitored daily for clinical appearance, and body weight and grip strength were measured every 4 days. Mice were euthanized when they were unable to move or were trembling. The experiment was terminated 28 days after the start of AD feeding, and survival was plotted as Kaplan–Meier curves.

### Tenotomy

To induce muscle hypertrophy, tenotomy was performed for the left leg of the mice as described previously (19). In anaesthetized mice, an incision was made along the midline of the calf following fur removal. Soleus and gastrocnemius (GAS) muscle tendons were excised, and then the skin was closed using three or four stitches. The right leg was subjected to sham surgery, in which the incision was immediately closed without tendon removal. Mice were killed, and the plantaris muscles were harvested following 14 days of overloading.

### Grip strength and tetanic muscle force

Whole-limb grip strength was measured using a grip strength meter for mice (Columbus Instruments). Peak tension (in N) was recorded when the mouse released its grip. The maximal strength determined with ten successive measurements was used for data analysis. Tetanic muscle force was measured in the tibialis anterior (TA) muscle using the Whole Animal Muscle Test System (Aurora Scientific) as previously described (19).

### Rotarod and treadmill running

Motor coordination was assessed using a rotarod apparatus (BioResearch Center, Japan) under a constant-speed protocol (6 rpm). The longest latency to fall among three separate trials was used for data analysis. Endurance capacity was evaluated using a treadmill running test. Mice were familiarized with the treadmill on the two days preceding the test (10 min at 10 m/min). On the test day, mice ran at 10 m/min on a 15° incline for 30 min, after which the speed was increased by 2 m/min every 15 min until exhaustion. Exhaustion was defined as the inability to resume running while remaining on the electrical stimulus grid at the back of the treadmill for more than 5 s. The total running distance was calculated and used for subsequent analyses.

### Myofiber isolation, culture, and diameter measurement

Individual myofibers were isolated from Extensor digitorum longus (EDL) muscles by enzymatic digestion with type I collagenase as previously described (19). Isolated myofibers were cultured for 48 h on Matrigel-coated plates in DMEM containing 10% horse serum supplemented with rDll4 (0.5, 1, 5, 50, or 500 ng/ml), rDll4-Fc (50 ng/ml), or untagged recombinant Dll1 protein (50 ng/ml; rDll1; 140-08, Thermo Fisher Scientific). For rescue experiments, recombinant Notch1-Fc chimera protein (500 ng/ml; rN1-ECD; #3647-TK, R&D Systems), recombinant Notch2-Fc chimera protein (500 ng/ml; rN2-ECD; #3735-NT, R&D Systems), or recombinant Notch3-Fc chimera protein (500 ng/ml; rN3-ECD; #1559-NT, R&D Systems) was added together with rDll4. Cultured myofibers were imaged using a microscope (IX83, Olympus), and myofiber diameters were measured in at least 20 individual myofibers using cellSens software (Olympus). Neuromuscular junctions (NMJs) were visualized using fluorescent dye-conjugated α-bungarotoxin (αBTX), and the percentage of fragmented NMJs was quantified to assess NMJ degeneration.

### Cell culture and *in vitro* assays

Human umbilical vein endothelial cells (HUVECs) were obtained from Lonza (C2519A, lot no. 20TL266553) and cultured on type I collagen-coated dishes in HuMedia-EG2 (KE-2150S, Kurabo) supplemented with the full growth supplement set (growth medium, GM) according to the manufacturer’s instructions. For culture supernatant analyses, cells were maintained in serum-free basal medium supplemented with the growth supplement set. HUVECs were treated with LPS (100 or 500 ng/ml), tumor necrosis factor-α (2 or 10 ng/ml), interleukin-1β (10 ng/ml), or cobalt chloride (100 μM) for 24 h, and culture supernatants were collected for analysis.

To examine the effects of rDll4 on endothelial senescence, HUVECs were maintained under four conditions: control, rDll4 (10 ng/ml), rDll4 plus αDll4 (100 ng/ml), and rDll4 plus rDll4-Fc (25 ng/ml). Cells were passaged every 3 days and maintained from passage 3 to passage 5 for a total of 9 days in GM. At passage 3 (day 3), passage 4 (day 6), and passage 5 (day 9), senescence was assessed by SPiDER-βGal staining according to the manufacturer’s instructions (SG02, Dojindo), and the proportion of SPiDER-βGal^+^ cells was quantified from four ×10 fields per sample. Total RNA was collected at passage 5 for gene expression analysis.

To reduce *DLL4* expression, HUVECs were transfected with 20 nM *DLL4*-targeting siRNA (SASI_Hs02_00352665, Sigma-Aldrich) or control siRNA using Lipofectamine RNAiMAX (Thermo Fisher Scientific) for 24 h. The medium was then replaced with the serum-free medium described above, and cells were cultured for an additional 24 h before collection of cell lysates and culture supernatants.

For live-cell imaging of Dll4 protein dynamics, HUVECs were transfected with a *DLL4-CTD-eGFP* expression plasmid using Lipofectamine LTX with Plus Reagents (Thermo Fisher Scientific) according to the manufacturer’s instructions and cultured in GM. Time-lapse fluorescence images were acquired using an EVOS imaging system (FL Auto, Thermo Fisher Scientific) at 10-min intervals over a 10-h period. Time-lapse movies were generated at a playback rate of 6 frames per second.

### RNA extraction and quantitative PCR with reverse transcription

Total RNA was extracted from GAS muscles, kidneys, livers, and cultured cells using ISOGEN II (Nippon Gene) and the RNeasy kit (Qiagen) according to the manufacturer’s instructions. cDNA was prepared using a ReverTra Ace kit with genomic DNA remover (TOYOBO). qPCR was performed using a THUNDERBIRD SYBR qPCR mix (TOYOBO) and a real-time PCR detection system (CFX96, Bio-Rad). Primer sequences are listed in Table S4. Expression levels were normalized to those of *Tbp* in mouse skeletal muscle, *Gapdh* in mouse kidney and liver, and *GAPDH* in HUVECs.

### Bulk RNA sequencing (RNA-seq)

Total RNA was extracted from GAS muscles, kidneys, and livers using ISOGEN II and the RNeasy kit. Library preparation was performed by Novogene (Beijing). Sequencing data were generated at a depth of >40 million reads per sample on an Illumina NovaSeq platform using paired-end 150-bp sequencing strategies and converted into FASTQ files. The FASTQ files were then mapped to the Mus musculus mm10 reference genomes and transcriptome (GENCODE vM23/Ensembl 98) (62, 63) using Strand NGS v.3.4 software (Strand Life Sciences), and gene-level read counts were obtained. Downstream transcriptomic analyses, including identification of differentially expressed genes (DEGs), principal component analysis (PCA), heatmap generation, and KEGG pathway enrichment analysis (22), were performed using RNASeqChef (64). DEGs were identified using the following thresholds: fold change > 1.2 (Fig. 1F) or > 1.5 (Figs. 1O and 4), false discovery rate (FDR) < 0.05. The base-mean cutoff was set to 0, with no additional filtering based on mean normalized read counts. Read counts were normalized by the median-of-ratios method implemented in DESeq2.

### Single-nucleus (sn) RNA-seq

Nuclei were isolated from frozen TA muscles from the AD, AGVeh, and AGrD4 groups (n = 2 per group) using the Chromium Nuclei Isolation Kit (10x Genomics). Briefly, frozen muscle tissues were minced, homogenized in Lysis Buffer, and passed through a filtration column. Debris was subsequently removed by centrifugation using the Debris Removal Buffer. The isolated nuclei were washed twice and resuspended in Resuspension Buffer. Nuclei were stained with PI, and their final concentrations were determined under a fluorescence microscope (Olympus IX83). Nuclei adjusted to the appropriate concentration were loaded onto the 10x Chromium system using the Chromium Next GEM Single Cell 3′ Reagent Kit v.3.1 (10x Genomics) according to the manufacturer’s instructions. Prepared libraries were sequenced on a NovaSeq X plus platform (Illumina) at Macrogen Japan.

Raw sequencing reads were processed using the Cell Ranger pipeline (v.7.2.0; 10x Genomics). Reads were aligned to the mm10 mouse reference genome (gex-mm10-2020-A) to generate a gene-barcode matrix of unique molecular identifier (UMI) counts. Sample aggregation was performed using the cellranger aggr command without read depth normalization. Downstream analyses were conducted using the Seurat R package (v.5.3.0) in R (v.4.5.0). To ensure high-quality data, cells were excluded from downstream analysis if they expressed fewer than 500 genes, more than 4,000 genes, or possessed a mitochondrial gene content exceeding 10%. Log-normalization and scaling were applied to the filtered dataset, which comprised 32,272 genes across 49,819 high-quality nuclei (AD1: 4,685 nuclei; AD2: 4,361 nuclei; AGVeh1: 8,848 nuclei; AGVeh2: 5,847 nuclei; AGrD4-1: 16,427 nuclei; AGrD4-2: 9,651 nuclei). Individual datasets were integrated using the IntegrateLayers function via the Reciprocal PCA (RPCA) method. Following integration, the data were projected into a two- dimensional space using Uniform Manifold Approximation and Projection (UMAP). Cell clusters were manually annotated through the expression of canonical cell-type markers. Final cell type annotations and coordinates were exported into the Loupe Cell Browser (10x Genomics), in which gene expression visualization was performed.

For selected gene expression analyses, nuclei were further classified based on predefined marker expression criteria, independently of the cluster-level annotations described above. Specifically, myofiber nuclei were identified by the expression of Ttn together with at least one of the myosin heavy chain genes (*Myh2*, *Myh4*, or *Myh1*) and further classified as type IIa, IIb, or IIx based on *Myh2*, *Myh4*, or *Myh1* expression, respectively. Other cell populations were identified based on the expression of specific marker genes, including *Pax7* (satellite cells), *Pecam1* (endothelial cells), *Rgs5* (pericytes), *Myh11* (smooth muscle cells), *Plin1* (adipocytes), *Dcn* (fibroblasts/FAPs), *Tnc* (tendon/MTZ cells), *Col22a1* (myotendinous junction cells), *Chrne* (NMJ-associated cells), and *Ptprc* (immune cells). A threshold of 0 was used, and nuclei with normalized expression values greater than 0 were considered marker-positive. We subset the Seurat object by the assigned cell type labels and experimental conditions, and generated dot plots for predefined gene sets using Seurat’s DotPlot (assay = RNA). Dot size represents the fraction of expressing cells, and dot color represents scaled average expression (default Seurat scaling). Plots were formatted with ggplot2.

### Histology and immunostaining

TA muscles and liver tissues were immediately frozen in liquid nitrogen-cooled 2-methylbutane and stored at −80°C until use. Frozen tissues were cryosectioned at 10 μm for TA muscles and 15 μm for liver tissues using a cryostat (Leica). For longitudinal TA sections, mice were perfusion-fixed with 4% paraformaldehyde (PFA). TA muscles were dissected, post-fixed in 4% PFA, cryoprotected in 30% sucrose, and cryosectioned longitudinally at 30 μm. Kidneys were fixed in 10% neutral buffered formalin, transferred to 70% ethanol, embedded in paraffin, and sectioned at 6 μm unless otherwise indicated. For retinal preparations, eyeballs were fixed in 4% PFA/PBS on ice for 30 min. Retinas were then isolated, briefly microwave-fixed in 4% PFA/PBS for 15 s on ice, and post-fixed in 4% PFA/PBS at 4°C overnight.

TA cryosections and isolated single myofibers were fixed in 4% PFA for 10 min, permeabilized with 0.3% Triton X-100/PBS for 20 min, blocked with 5% normal serum/PBS for 30 min, and incubated overnight at 4 °C with primary antibodies against laminin α-2 (1:600; sc-59854, clone 4H8-2, Santa Cruz), MyHC IIa (1:3; clone SC-71, DSHB), MyHC IIb (1:3; clone BF-F3, DSHB), CD31 (1:400; 102502, clone MEC13.3, BioLegend), dystrophin (1:400; ab15277, abcam), and Pax7 (1:200; sc-81648, clone PAX7, Santa Cruz). Samples were then incubated for 1 h at room temperature (20–25°C) with Alexa Fluor–conjugated secondary antibodies: goat anti-rat IgG Alexa Fluor 488 (1:500; A11006, Thermo Fisher Scientific), goat anti-rat IgG Alexa Fluor 546 (1:500; A11081, Thermo Fisher Scientific), donkey anti-rat IgG Alexa Fluor 555 (1:500; A48270, Thermo Fisher Scientific), goat anti-mouse IgM Alexa Fluor 488 (1:500; AB150121, abcam), goat anti-mouse IgG1 Alexa Fluor 594 (1:500; A21125, Thermo Fisher Scientific), donkey anti-rabbit IgG Alexa Fluor 647 (1:500; A31573, Thermo Fisher Scientific), and goat anti-mouse IgG Alexa Fluor 488 (1:500; A11029, Thermo Fisher Scientific). For SPiDER-βGal staining of TA cross-sections, sections were first stained with SPiDER-βGal according to the manufacturer’s instructions (SG02, Dojindo), followed by secondary antibody incubation. The number of SPiDER-βGal–positive cells were quantified from 4–5 fields per section acquired at ×20 magnification. Myofiber cross-sectional area was quantified either from whole TA cross-sections or from four ×10 fields in the outer region of the section, depending on the experiment, using Fiji/ImageJ or AIVIA. Vascular area in TA sections was quantified from three 500 × 500 μm fields per section using Fiji/ImageJ.

For longitudinal TA sections, sections were blocked with normal donkey serum for 1 h at room temperature, incubated overnight at 4°C with primary antibodies against Dll4 (1:100; sc-365429, clone G-12, Santa Cruz), dystrophin, CD31, collagen I (1:500; 1310-01, SouthernBiotech), and collagen IV (1:500; 1340-01, SouthernBiotech), and then incubated overnight at 4°C with Alexa Fluor–conjugated secondary antibodies: donkey anti-mouse IgG Alexa Fluor 488 (1:500; A21208, Thermo Fisher Scientific), donkey anti-rabbit IgG Alexa Fluor 405 (1:500; A48258, Thermo Fisher Scientific), donkey anti-rat IgG Alexa Fluor 555, donkey anti-goat IgG Alexa Fluor 546 (1:500; A11056, Thermo Fisher Scientific), and donkey anti-goat IgG Alexa Fluor 647 (1:500; A21447, Thermo Fisher Scientific). In collagen I staining experiments, sections were fixed in ice-cold acetone at −80°C for 15 min before immunostaining.

For retinal vascular staining, retinal preparations were incubated with anti-CD31 antibody, followed by overnight incubation at 4°C with Alexa Fluor 555–conjugated secondary antibody. Retinal vascular images were acquired with an Andor BC43 benchtop confocal microscope, and vascular density was quantified from three 500 × 500 μm fields per retina using Fiji/ImageJ.

Liver sections were stained with SPiDER-βGal according to the manufacturer’s instructions (Dojindo). The number of SPiDER-βGal–positive cells were quantified from five fields per section acquired at ×20 magnification using Fiji/ImageJ.

For kidney immunostaining, deparaffinized and rehydrated sections were subjected to antigen retrieval in Dako antigen retrieval buffer (S1699, Agilent) using an autoclave. Sections were blocked with 5% normal donkey serum/PBS, incubated overnight at 4°C with primary antibodies against Dll4, CD34 (1:400; 553738, clone RAM34, BD Biosciences), and collagen IV, and then incubated with Alexa Fluor–conjugated secondary antibodies for 1 h at room temperature. Glomerular vascular area was quantified as the CD34-positive area normalized to glomerular area in at least 12 glomeruli per section, and Dll4 staining intensity was quantified from six fields per section acquired at ×40 magnification, using Fiji/ImageJ.

For renal histology, Masson’s trichrome and periodic acid–Schiff (PAS) staining were performed on paraffin-embedded kidney sections. Masson’s trichrome staining was performed on 2-μm sections using a Masson’s Trichrome Stain kit (TRG-1, ScyTek Laboratories), and PAS staining was performed on 6-μm sections using a PAS Stain kit (PAS-1, ScyTek Laboratories), according to the manufacturer’s instructions. Fibrotic area in Masson’s trichrome-stained sections was quantified from four ×10 fields per section using Fiji/ImageJ. Bowman’s capsule length was measured in at least 20 glomeruli per section as previously described (65). Tubular injury was assessed in four ×10 fields per section by counting tubules exhibiting protein cast formation, tubular dilatation, vacuolization, and tubular cell degeneration. The total number of affected tubules was used as the tubular injury score.

For immunofluorescence staining of cultured HUVECs, cells were then fixed with 2% PFA in PBS for 5 min and blocked with 5% normal goat serum in PBS for 30 min without permeabilization. Cells were incubated overnight at 4°C with a rabbit antibody against the N-terminal domain of Dll4 (1:400; 500-P279, PeproTech/Invitrogen), followed by goat anti-rabbit IgG Alexa Fluor 488 (1:800; A-11008, Thermo Fisher Scientific) or Alexa Fluor 555 (1:800; A-21429, Thermo Fisher Scientific) for 1 h at room temperature. For WGA and Dll4 co-staining, living cells were incubated with Alexa Fluor 594– conjugated wheat germ agglutinin (WGA; 1:500; W11262, Thermo Fisher Scientific) in PBS for 10 min in a CO₂ incubator before fixation.

Images were acquired using an Olympus IX83 fluorescence microscope, an Olympus FV4000 confocal microscope, or an Andor BC43 benchtop confocal microscope for retinal vascular imaging. Quantitative image analyses were performed using Fiji/ImageJ or AIVIA (Leica).

### Proximity ligation assay

In situ proximity ligation assay (PLA) was performed on 30-µm longitudinal cryosections of perfusion-fixed mouse TA muscles using the Duolink In Situ Red Starter Kit Mouse/Goat (DUO92103, Sigma-Aldrich) according to the manufacturer’s instructions, with minor modifications. Sections were washed with PBS and permeabilized with 0.3% Triton X-100 in PBS for 15 min at room temperature. After blocking with Duolink Blocking Solution for 1 h at 37°C, sections were incubated overnight at 4°C with mouse anti-Dll4 (1:100; sc-365429, clone G-12, Santa Cruz) and goat anti-collagen I (1:500; 1310-01, SouthernBiotech) primary antibodies diluted in Duolink Antibody Diluent. After washing with Wash Buffer A, sections were incubated with anti-mouse PLUS and anti-goat MINUS PLA probes for 90 min at 37°C. Ligation and rolling circle amplification were then performed using the Duolink ligation and amplification reagents. PLA signals were detected using the Red detection reagent. Nuclei were counterstained with DAPI, and transmitted light images were acquired to visualize tissue morphology. Images were acquired by confocal microscopy (Olympus FV4000) using identical acquisition settings and the same Z-depth range for all samples. Projection images were used for quantification. PLA puncta were quantified using Fiji/ImageJ from the red channel with a fixed intensity threshold applied uniformly to all images. Puncta were counted within a fixed imaging area of 320 × 320 µm² and expressed as PLA puncta per field. No punctate signals were observed in no-primary control sections.

### Immunoelectron microscopy

Under deep anesthesia by inhalation of isoflurane, aged mice (26-month-old) were fixed by perfusion through the ascending aorta with 4% PFA and 0.1% glutaraldehyde in 0.1 M phosphate buffer (PB). Serial 70-μm-thick sections were cut using a vibrating microtome (Dosaka TTK-3000) from the GAS muscles and processed for immunogold labeling. After cryoprotection with 25% sucrose in PB and rapid freeze-thawing in the vapor of liquid nitrogen, sections were incubated with anti-Dll4 antibody (1:200) in 0.01M phosphate buffered saline containing 5% donkey serum at 4 °C for 7 d, followed by incubation with a 10 nm colloidal gold-labelled goat anti-mouse secondary antibody (BBI Solutions) overnight. Sections were then post-fixed with 1% OsO₄, stained en bloc with 1.5% uranyl acetate, dehydrated, subjected to a substitution process, and embedded in epoxy resin. Ultrathin sections were cut using an ultramicrotome (Leica EM UC7) and then stained with uranium acetate and lead citrate. Images were captured using a transmission electron microscope (Hitachi H7700). Consistent immunogold labeling patterns were observed in samples from three independent mice.

### NAD⁺ levels

Hepatic NAD⁺ levels were measured by high-performance liquid chromatography (HPLC) as previously described (66). Briefly, liver tissues were snap-frozen in liquid nitrogen and homogenized in 10% perchloric acid. The resulting supernatants were neutralized with 3 M K₂CO₃, and NAD⁺ levels were determined using a Shimadzu HPLC system equipped with a SUPELCOSIL LC-18-T column (Merck). NAD⁺ levels were normalized to tissue weight.

### Cell-free extracellular matrix (ECM)-binding assay

rDll4 and rDll4-Fc (10185-D4) were used in the cell-free ECM-binding assay. Briefly, 2 µg/ml of recombinant proteins were incubated in the wells coated with type I collagen (354249, BD biosciences), type IV collagen (C5533, Sigma-Aldrich), and Matrigel (rich in laminin, collagen IV, entactin/nidogen, and heparan sulphate proteoglycans; 356231, Corning) at 37°C for 2 h. All ECM substrates were used at a coating concentration of 1 mg/ml. After incubation, the supernatants were collected, and the remaining proteins were analyzed by immunoblotting. An aliquot of the protein solution equivalent to the volume added to each well was immediately mixed with SDS sample buffer and boiled to serve as an input control.

### Protein extraction and immunoblotting

Total protein lysates were obtained from homogenized tissues and cultured cells using RIPA buffer (FUJIFILM). HUVEC culture supernatants were cleared by sequential centrifugation––500 × g for 10 min, 12,500 × g for 40 min at 4°C, and then ultracentrifugation at 100,000 × g for 2 h at 4°C––to remove dead cells and extracellular vesicles. The resulting supernatants were collected and concentrated using Amicon Ultra (UFC900324, Merck) to quantify the soluble form of proteins in the culture media. Protein concentration in each sample was quantified using a Pierce BCA Protein Assay Kit (Thermo Fisher Scientific). Primary antibodies were diluted in CanGetSignal solution A (TOYOBO) or 5% skim milk and incubated with membranes at 4 °C overnight. Primary antibodies included N- terminal anti-Dll4 antibodies for mouse and human Dll4 detection (1:1,000; 21584-1-AP, proteintech), a C-terminal anti-Dll4 antibody for human Dll4 detection (1:1,000; #96406, clone D7N3H, Cell Signaling Technology), and an antibody against β-tubulin (1:10,000; 010-25043, clone 10G10, FUJIFILM). Membranes were then washed and incubated for 1 h at room temperature with horseradish peroxidase (HRP)-conjugated secondary antibodies diluted in CanGetSignal solution B (TOYOBO) or 5% skim milk: HRP-conjugated anti-rabbit IgG (1:10,000; #7074, Cell Signaling Technology) and HRP-conjugated anti-mouse IgG (1:10,000; #7076, Cell Signaling Technology). Secondary antibodies were visualized using chemiluminescence with a LAS-4000 digital luminescent image analyzer (GE Healthcare) or a FUSION western blot imaging system (Vilber). Immunodetection of β-tubulin and ponceau staining served as internal controls.

### Immunoprecipitation

Protein lysates from GAS muscles and HUVECs were prepared in immunoprecipitation (IP) lysis buffer (Thermo Fisher Scientific). GAS lysates, HUVEC lysates, and concentrated HUVEC culture supernatants were incubated for 20 min at room temperature with anti-Dll4 antibody–conjugated Dynabeads (Thermo Fisher Scientific), prepared by cross-linking anti-Dll4 antibodies to Dynabeads using dimethyl pimelimidate (DMP) (sc-365429, Santa Cruz, for GAS samples; #96406, CST, for HUVEC samples). Bead-bound immune complexes were collected using a magnetic separator and washed at least five times with 0.025% Tween-20/PBS. Proteins associated with the antibody-conjugated beads were eluted by incubating washed beads in IP elution buffer (10 mM Tris-HCl, 300 mM NaCl, 5 mM EDTA and 0.5% SDS) for 10 min at 70 °C. Eluted proteins were used for either immunoblotting or liquid chromatography–tandem mass spectrometry (LC–MS/MS) analysis.

For IP-immunoblotting, eluted proteins from GAS samples were separated by SDS-PAGE and transferred to membranes. Membranes were blocked with 5% skim milk and probed with antibodies against Dll4 (sc-365429, Santa Cruz) and collagen I (1:1,000; 1310-01, Southern Biotech), followed by HRP-conjugated secondary antibodies: anti-mouse IgG-HRP and anti-goat IgG-HRP (1:10,000; sc-2056, Santa Cruz). Non-immunoprecipitated input lysates were used as loading controls.

### LC–MS/MS analysis

Immunoprecipitated proteins were used for LC–MS/MS analysis. Eluted proteins were separated by SDS-PAGE and visualized using a silver staining kit (24600, Thermo Fisher Scientific). Bands of interest were excised from the gel based on the molecular weight marker and minced into 1 mm^3^ pieces. Destaining was performed according to the manufacturer’s instructions, followed by reduction for 10 min at 60 °C using dithiothreitol and carbamidomethylation for 30 min at room temperature using iodoacetamide. After dehydration, in-gel digestion was performed at 25 °C for 17 h using Glu-C endoproteinase sequencing grade from *Staphylococcus aureus* V8 (150 ng/band; Roche Diagnostics). The resulting peptides extracted from each band were desalted using GL-tip SDB (GL Sciences), and the eluates were concentrated in vacuo, dissolved in 5 % MeCN containing 0.1 % formic acid (FA), and subjected to LC–MS/MS analysis.

LC–MS/MS analyses were performed on a Vanquish Neo (Thermo Fisher Scientific) coupled to an Orbitrap Fusion Lumos mass spectrometer (Thermo Fisher Scientific) equipped with an electron spray ionization source. The LC system was equipped with a trap column (C18 PepMap Neo, 0.3 × 5 mm, 5 μm, Thermo Fisher Scientific) and an analytical column (NTCC-360/ 75–3-125, Nikkyo Technos). Peptide separation was performed using a 65-min gradient of water/0.1% FA (mobile phase A) and MeCN/0.1% FA (mobile phase B) at a flow rate of 300 nL/min. Elution was performed as follows: 0–1 min, 5 % B; 1–61 min, 5–22 % B; 61–66 min, 22–40 % B; 66–70 min, 40–90 % B; and 70–75 min, 90 % B. The mass spectrometer was operated in data-dependent acquisition mode. The MS parameters were as follows: spray voltage, 2.0 kV; capillary temperature, 275 °C; S-lens RF level, 30; scan type, full MS; scan range, *m/z* 350–1500; resolution, 60,000; polarity, positive; automatic gain control target, standard; and maximum injection time, auto. The MS/MS parameters were as follows: resolution, 15,000; automatic gain control target, standard; maximum injection time, auto; normalized collision energy (NCE), 30; dynamic exclusion, 20 s; isolation window, 1.6 *m/z*; and included charge states, 2–7.

The identification of proteins was performed using Proteome Discoverer software ver 3.2 (Thermo Fisher Scientific). The analytical parameters used for the database search were as follows: parent mass tolerance, 10.0 ppm; fragment mass tolerance, 0.02 Da; search engine, Sequest HT; protein database, DLL4_HUMAN (Uniprot); enzyme name, GluC (full + semi); maximum number of missed cleavages, 2; dynamic modification, oxidation (methionine), acetyl (lysine), phospho (serine, threonine, tyrosine), acetyl (N-terminus), Met-loss (protein terminus methionine), and Met-loss+acetyl (protein terminus methionine); static modification, carbamidomethyl (cysteine); and confidence, FDR < 0.01.

### ELISA

Mouse whole blood was collected from the retro-orbital plexus under anesthesia without anticoagulant, allowed to clot at room temperature for 60 min, and centrifuged at 3,000 × g for 10 min at 4°C. Mouse urine samples were collected and centrifuged at 3,000 × g for 10 min at 4°C to remove debris. Human peripheral blood was drawn into serum-separating tubes, allowed to clot at room temperature for 30– 60 min, and centrifuged under the same conditions. The resulting serum was carefully collected and stored at −80°C until analysis. Dll4 levels in mouse serum, mouse urine, and human serum were measured using species-specific sandwich ELISA kits (High sensitivity Mouse Dll4 ELISA kit, SK00538-03, Aviscera Bioscience; Human Dll4 ELISA kit, SK00538-01, Aviscera Bioscience) according to the manufacturer’s instructions with minor modifications. Briefly, standards, serum samples (diluted fourfold in the kit dilution buffer), and urine samples were added to pre-coated 96-well plates (100 µl/well) and incubated for 2 h on the plate shaker at room temperature. After washing, HRP-conjugated detection antibody was added and incubated for 60–90 min with protection from light. Plates were developed with TMB substrate for 15–30 min, and the reaction was stopped using the stop solution. Absorbance at 450 nm was measured using a microplate reader, and analyte concentrations were calculated from the standard curve.

### Human cohort study

Data were obtained from the Arao cohort, a subset of the Japan Prospective Studies Collaboration for Aging and Dementia (JPSC-AD), a population-based prospective cohort study of community-dwelling adults aged 65 years or older conducted at eight sites in Japan (67, 68). The present study analyzed baseline data from 1,577 participants recruited in Arao City, Kumamoto Prefecture, Japan, between 2016 and 2017. After excluding 128 participants with missing covariate data, 1,449 participants were included in the analysis. The study protocol was approved by the Ethics Committee of Kumamoto University, for the parent cohort study (GENOME-333) and for the present analysis using existing specimens and data (No. 3099). The study was conducted in accordance with the Declaration of Helsinki. Written informed consent was obtained from all participants at the baseline survey. The present analysis was observational and involved no intervention.

### Statistical analysis for animal and cell experiments

No statistical method was used to predetermine sample sizes for animal and cell experiments. Sample sizes were selected based on previous studies using similar experimental approaches and are indicated in the figure legends. Statistical analyses of animal and cell experiments were performed using GraphPad Prism 11. Comparisons between two groups were performed using unpaired two-tailed Student’s t-tests, while comparisons among multiple groups were performed using one-way or two-way ANOVA followed by Tukey’s multiple comparisons test, unless otherwise indicated. Exceptions were as follows: Figure 2P, two-way ANOVA followed by Šídák’s multiple comparisons test, with an additional one-sample t-test against 100% input; Figure 4Q, mixed-effects model (REML) with group and time as fixed effects, individual mice as a random effect, and a group × time interaction, followed by Šídák’s multiple comparisons test; Figures 4R and 6P, log-rank test. Data are presented as mean ± SEM unless otherwise indicated. A P value < 0.05 was considered statistically significant. Statistical details are provided in the figure legends.

### Statistical analysis for human study

Statistical analyses of the human cohort were performed using IBM SPSS Statistics (version 29; IBM). Estimated glomerular filtration rate (eGFR) was calculated from serum creatinine, age, and sex using the equation proposed by the Japanese Society of Nephrology (eGFR = 194 × Cr^-1.094^ × Age^-0.287^ (male); eGFR = 194 × Cr^-1.094^ × Age^-0.287^ × 0.739 (female), where Cr represents serum creatinine (mg/dL) and age is expressed in years). Kidney function was categorized as G1 (≥90), G2 (60–89), G3a (45–59), G3b (30–44), and G4–G5 (<30 mL/min/1.73 m²). Reduced kidney function was defined as eGFR <60 mL/min/1.73 m².

Because serum Dll4 levels were not normally distributed, nonparametric tests were used. Differences in serum Dll4 concentrations across age categories and kidney function stages were assessed using the Kruskal–Wallis test followed by Dunn’s multiple comparisons test. Associations between serum Dll4 levels and clinical variables were evaluated using Spearman’s rank correlation analysis. Participant characteristics across serum Dll4 quintiles were compared using the Kruskal– Wallis test for continuous variables and the χ² test for categorical variables. Logistic regression analyses were performed to estimate odds ratios and 95% confidence intervals for reduced kidney function across serum Dll4 quintiles, using the lowest quintile as the reference and sequentially adjusting for potential confounders. Continuous variables are presented as medians and interquartile ranges, and categorical variables as percentages, unless otherwise indicated. Human cohort data in Figures 6A and 6B are presented as Tukey box plots.

## Data availability

The RNA-seq data have been deposited in the Gene Expression Omnibus under the following accession codes: snRNA-Seq, GSE329242; bulk RNA-Seq for Fig. 1F, GSE329674; bulk RNA-Seq for Fig. 1O, GSE329319; liver bulk RNA-Seq for Fig. 4H–I, GSE329581; and kidney bulk RNA-Seq for Fig. 4K–L, GSE329517. Other raw data are available upon reasonable request to the corresponding author.

## Acknowledgments

We thank S. Burden for providing *Mlc1f-Cre* mice; R. Nishinakamura and S. Tanigawa for their technical advice; and all members of the Ono laboratory for their helpful comments.

## Author contributions

ShF conducted and performed the experiments, interpreted the data, assembled the input data, and wrote the manuscript. NH, RO, HPOQ, KN, KS, SI, YoM, and YG performed cell-culture and animal experiments. TM provided key materials and critical comments. YT, YutaM, and TF performed the electron microscopy and immunohistochemical experiments, and wrote the manuscript. SaF performed histochemical experiments. KY and SU performed NGS analysis and wrote the manuscript. DA, NK, and MT analyzed human cohorts, interpreted the data, and wrote the manuscript. Yuri M performed the LC–MS/MS analysis, interpreted the data, and wrote the manuscript. TE and NI measured the NAD levels, interpreted the data, and wrote the manuscript. YO designed and performed the experiments, interpreted the data, assembled the input data, and wrote the manuscript. All authors reviewed and approved the final manuscript.

---

**Fig. S1.**
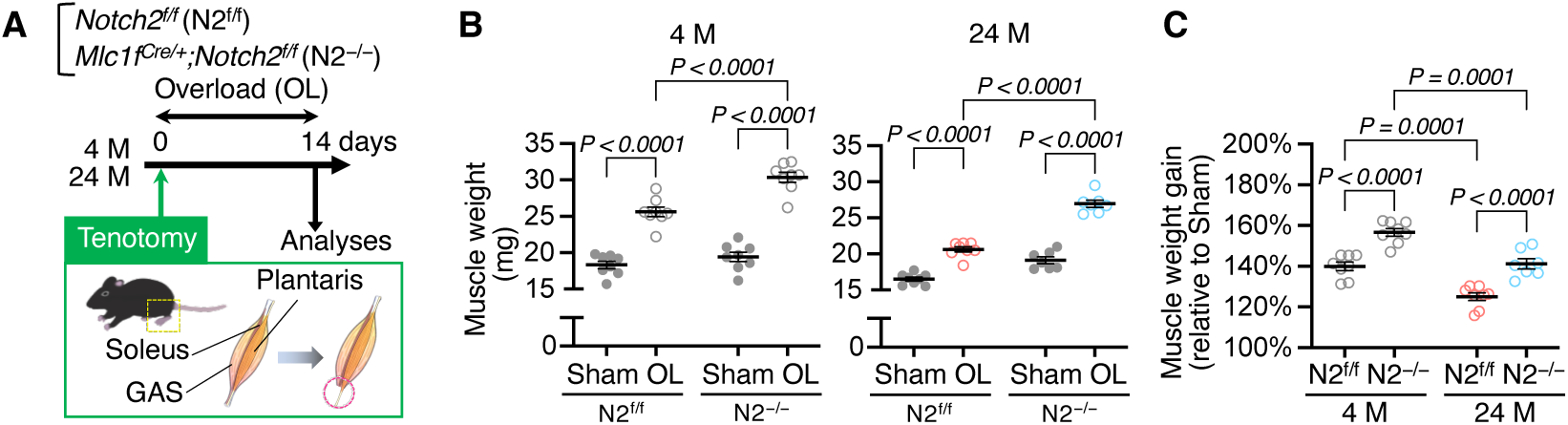
Notch2 inactivation alleviates age-related anabolic resistance. A–C, N2^f/f^ and N2^−/−^ mice aged 4 or 24 months were subjected to tenotomy-induced functional overload. Plantaris muscles were collected 2 weeks after surgery. **A,** Experimental scheme. **B,C,** Plantaris muscle weights after 2 weeks of overloading (**B**) and overload-induced weight gain relative to sham (**C**) (n = 7–8). Data are shown as mean ±SEM. P values were determined by two-way ANOVA with Tukey’s post-hoc test.

**Fig. S2.**
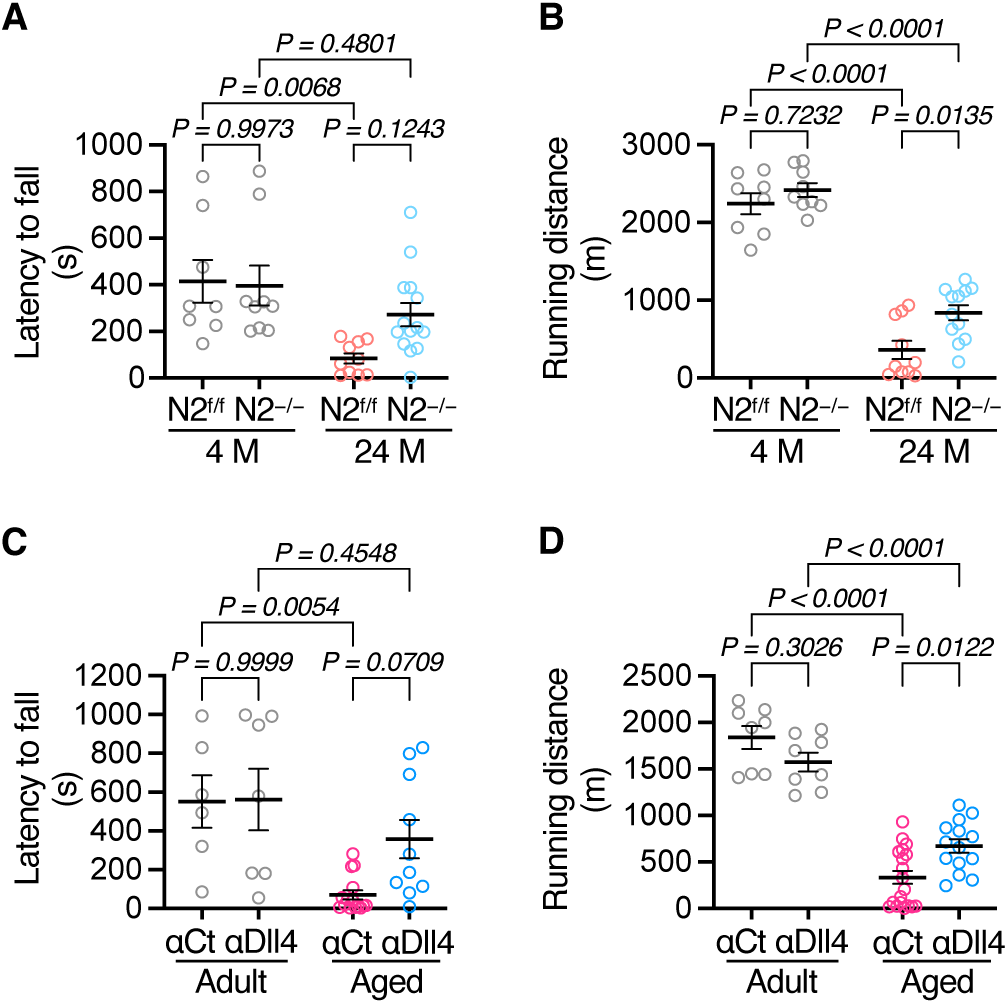
Dll4–Notch2 axis blockade attenuates age-related decline in physical performance. A,B, Rotarod latency to fall (**A**) and treadmill running distance (**B**) in N2^f/f^ and N2^−/−^ mice at 4 or 24 months of age (4 M and 24 M; n = 8–9 for 4 M; n = 10–14 for 24 M). **C,D**, Rotarod latency to fall (**C**) and treadmill running distance (**D**) in wild-type mice treated with anti-Dll4 neutralizing antibody (αDll4; 0.5 mg/kg) or control IgG (αCt) every 4 days for 4 months, starting at 3 (adult) or 21 (aged) months of age (n = 6–8 for adult; n = 10–20 for aged). Data are shown as mean ±SEM. P values were determined by two-way ANOVA with Tukey’s post-hoc test.

**Fig. S3.**
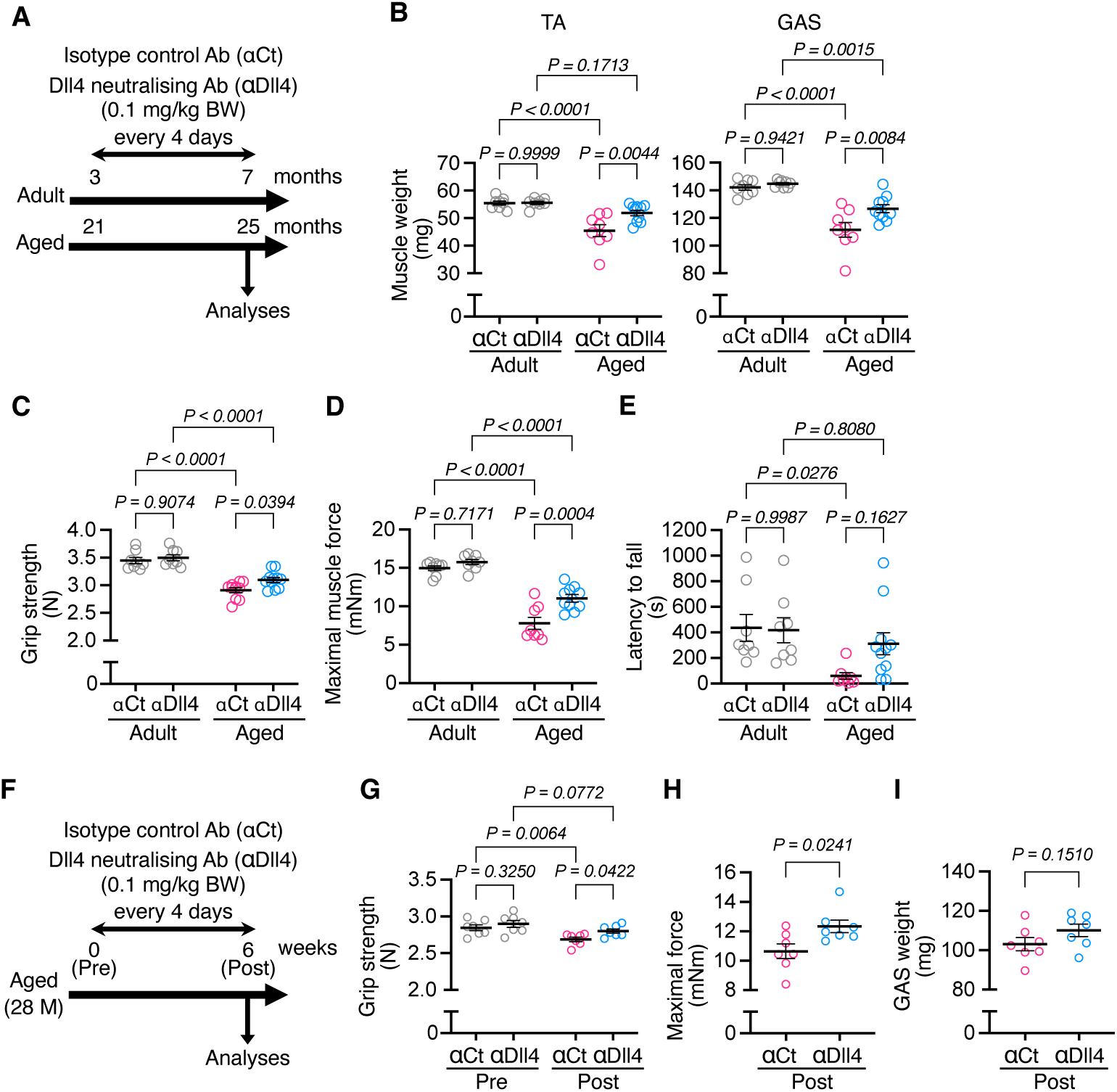
Low-dose αDll4 treatment preserves muscle mass and function in aged and geriatric mice. A–E, Wild-type mice were administered αDll4 (0.1 mg/kg) every 4 days for 4 months from 3 or 21 months of age; αCt was used as a control. **A**, Experimental scheme. **B–E**, Tibialis anterior (TA) and gastrocnemius (GAS) muscle weights (**B**), grip strength (**C**), TA maximal muscle force (**D**), and rotarod latency to fall (**E**) (n = 8–11). **F**, Experimental scheme: 28-month-old wild-type mice were injected with αDll4 (0.1 mg/kg) every 4 days for 6 weeks; αCt served as a control. (**G–I**) Grip strength (**G**), TA maximal muscle force (**H**), and GAS muscle weight (**I**) (n = 7). Data are shown as mean ±SEM. P values were determined by two-way ANOVA with Tukey’s post hoc test (**B–E,G**), and Student’s two-tailed unpaired t-test (**H,I**).

**Fig. S4.**
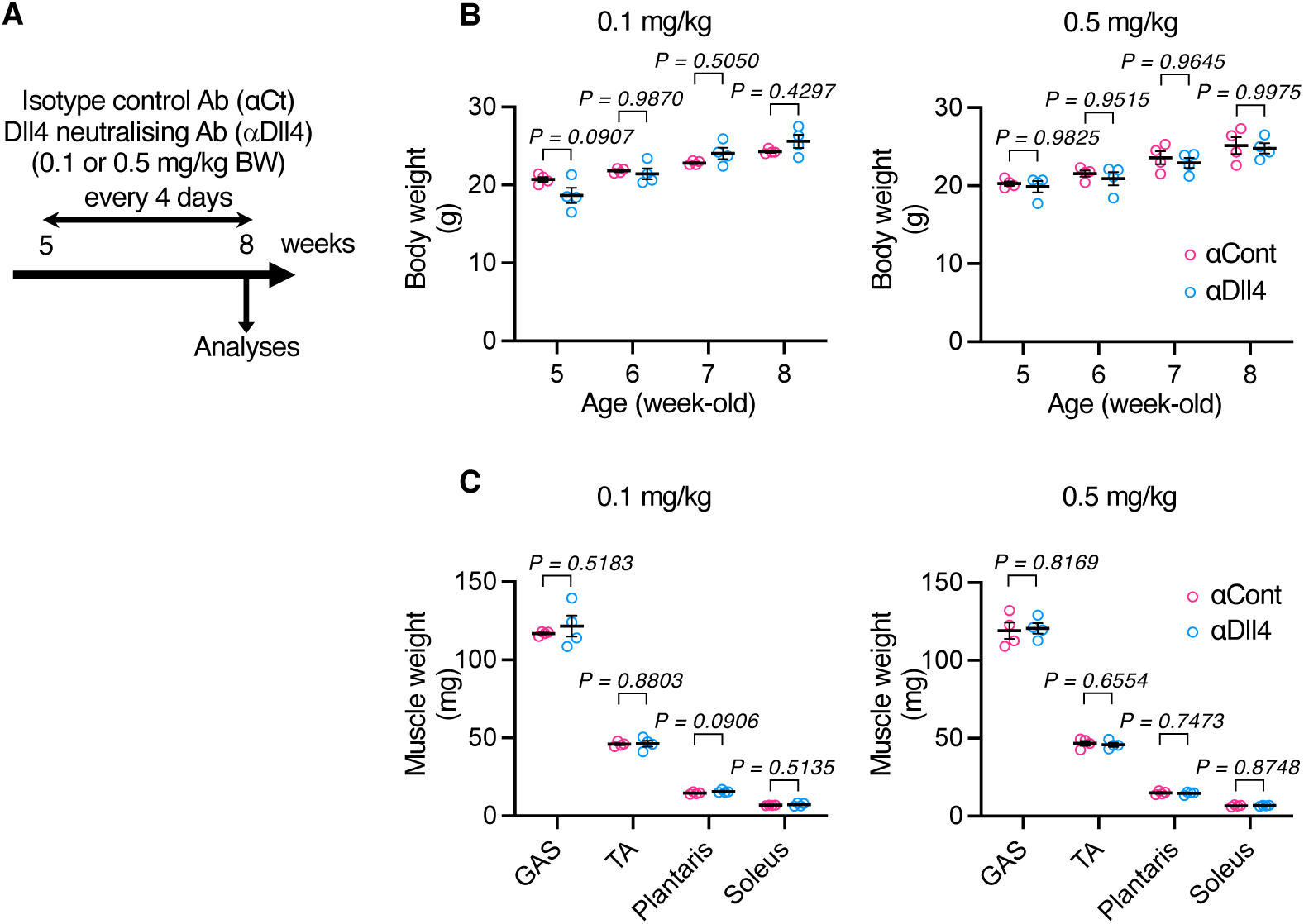
αDll4 treatment does not affect mouse growth. A–C, Wild-type mice were administered αDll4 (0.1 or 0.5 mg/kg) every 4 days for 3 weeks from 5 or 8 weeks of age; αCt was used as a control. **A**, Experimental scheme. **B**, Changes in body weight. **C**, Weights of GAS, TA, plantaris, and soleus muscles (n = 4). Data are shown as mean ±SEM. P values were determined by repeated measures ANOVA with Šídák’s post-hoc test (**B**), and Student’s two-tailed unpaired t-test (**C**).

**Fig. S5.**
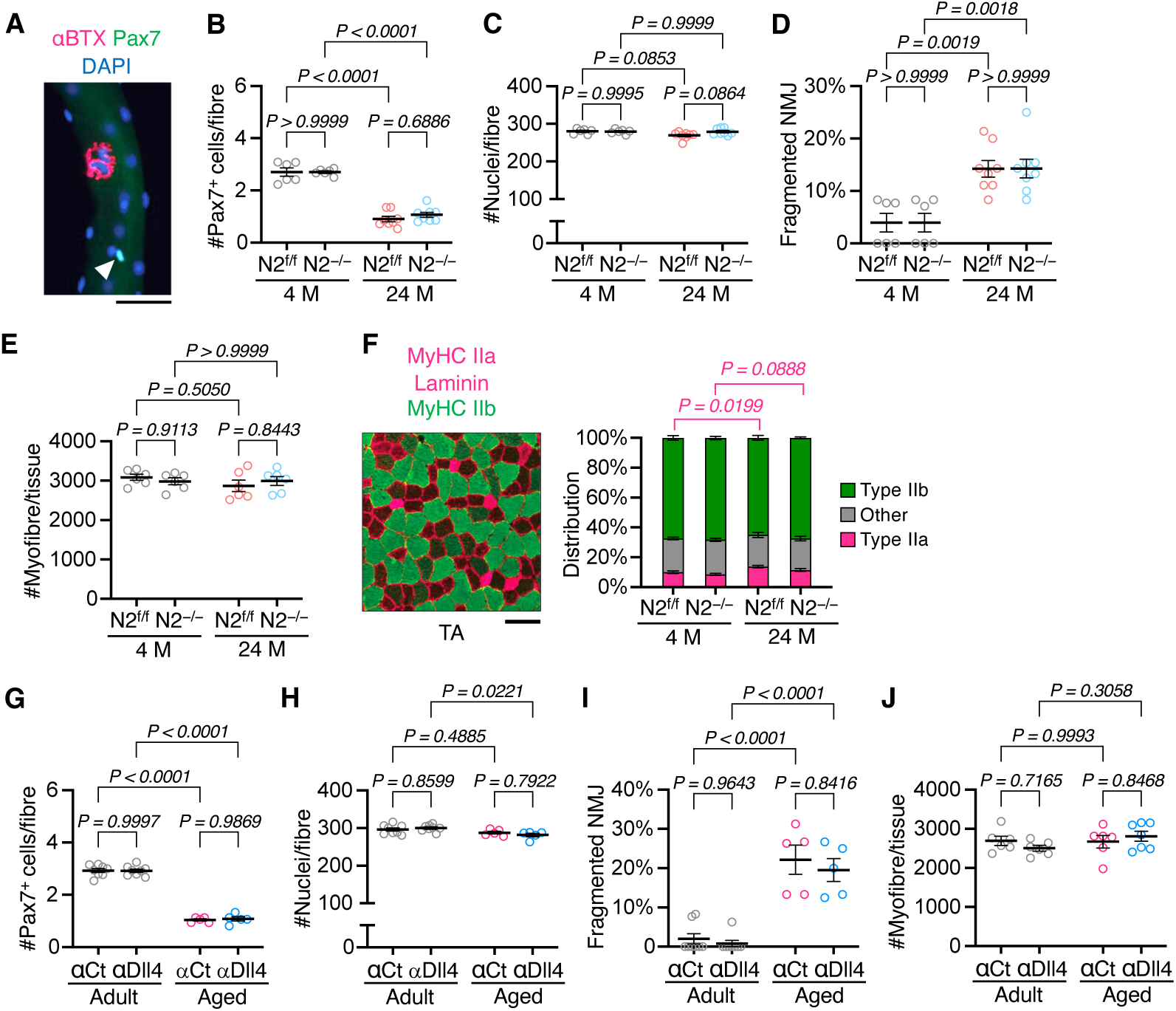
Dll4–Notch2 blockade does not affect satellite cells, myonuclei, myofiber composition, and neuromuscular junctions. A, Representative image of an isolated myofiber stained for Pax7 (green), α-bungarotoxin (αBTX; magenta), and DAPI (scale bar, 50 µm). Arrowhead indicates a satellite cell. **B–E**, Satellite cell and myonuclear numbers per myofiber (**B,C**), percentage of fragmented neuromuscular junctions (NMJs) (**D**), and myofiber number in TA muscles (**E**) from N2^f/f^ and N2^−/−^ mice at 4 and 24 months of age (4 M and 24 M; n = 8), as described in Fig. 1. **F**, Representative TA muscle sections stained for MyHC IIb (green), MyHC IIa/laminin (magenta), and quantification of myofiber-type distribution in N2^f/f^ and N2^−/−^ mice at 4 and 24 months (n = 6). **G–J**, Satellite cell number myonuclear numbers per myofiber (**G,H**), percentage of fragmented NMJs (**I**), and total myofiber number in TA muscles (**J**) from adult and aged mice treated with αDll4 or αCt (n = 5–8), as described in Fig. 1. Data are shown as mean ±SEM. P values were determined by two-way ANOVA with Tukey’s post-hoc test.

**Fig. S6.**
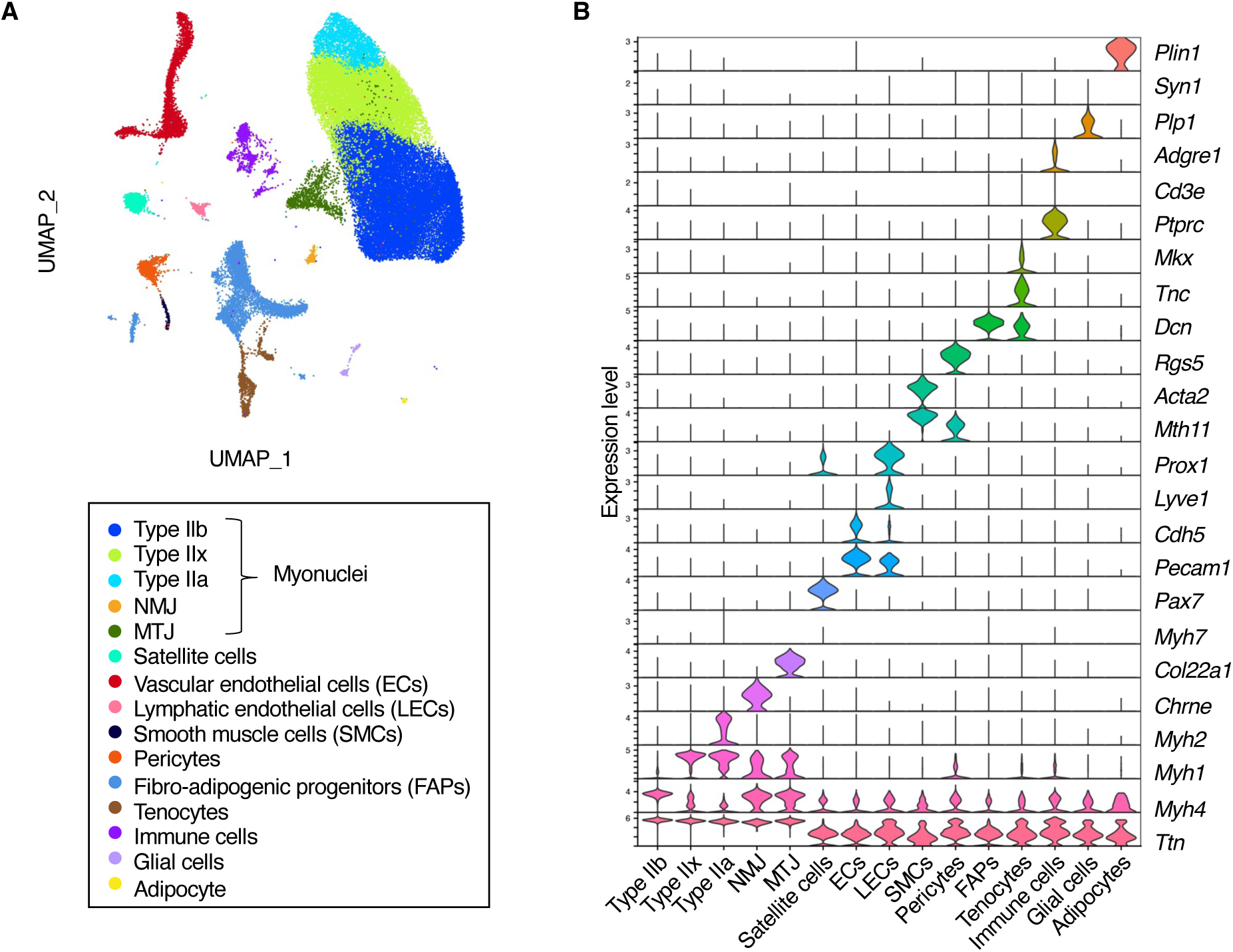
snRNA-seq analysis of adult and aged muscles following rDll4-Fc treatment. A, Uniform Manifold Approximation and Projection (UMAP) plot from unbiased clustering of 49,819 single nuclei isolated from TA muscles of adult mice (Adult) and aged mice treated with vehicle (Aged Veh) or rDll4-Fc (5 µg/kg; Aged rDll4-Fc). **B**, Violin plots showing the expression of representative marker genes across cell clusters identified by snRNA-seq.

**Fig. S7.**
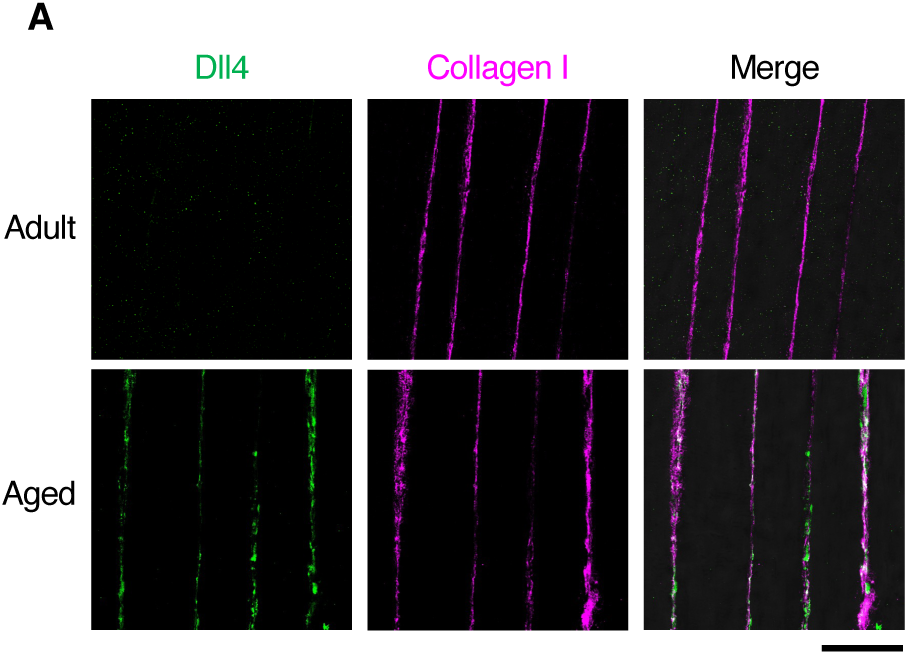
Dll4 colocalizes with collagen I in aged skeletal muscle. A, Representative longitudinal TA sections from adult (3-month-old) and aged (26-month-old) mice stained for Dll4 (*green*) and collagen I (*magenta*) (scale bar, 100 µm).

**Fig. S8.**
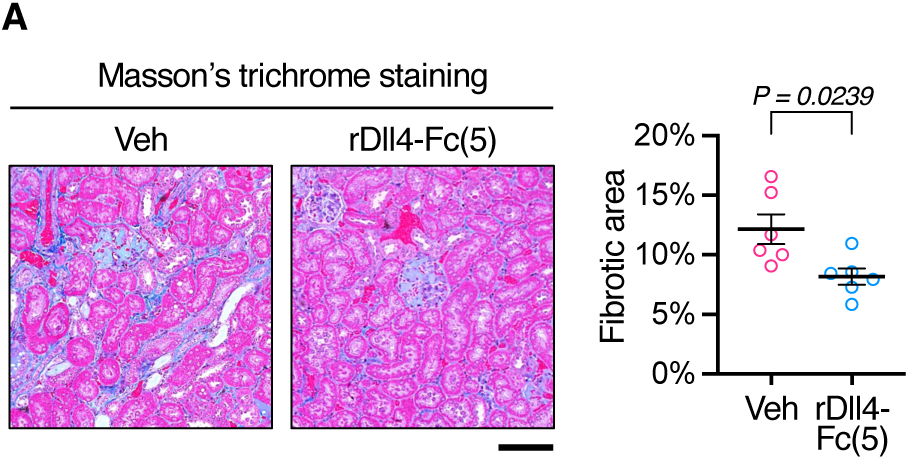
rDll4-Fc treatment alleviates age-related renal fibrosis. A, Representative Masson’s trichrome-stained kidney sections of aged mice treated with rDll4-Fc (5 µg/kg) or vehicle, with quantification of fibrotic area (n = 6; scale bar, 100 µm). Data are shown as mean ±SEM. P values were determined by Student’s two-tailed unpaired t-test.

**Fig. S9.**
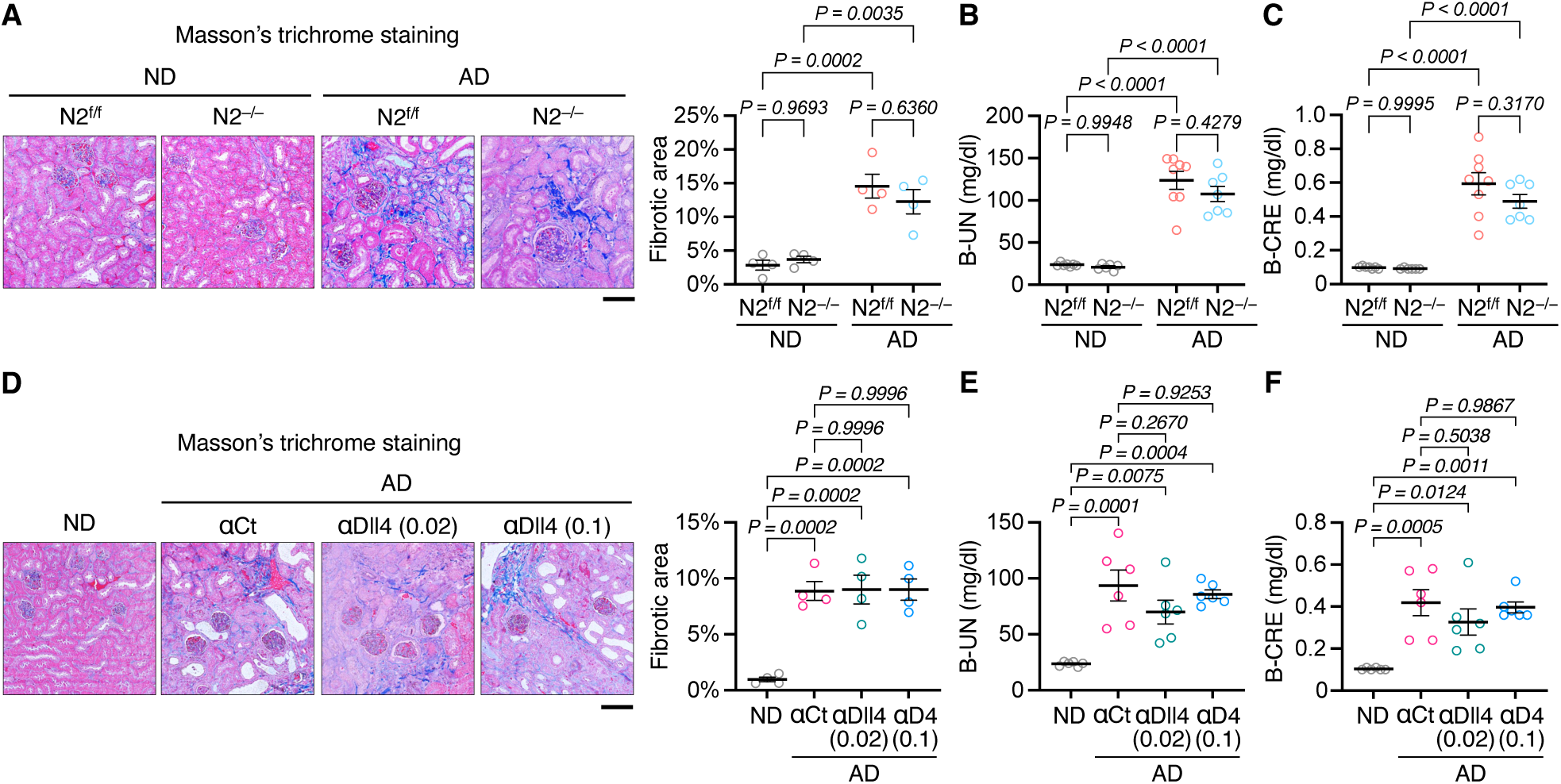
Characterization of chronic kidney disease mouse models. A–C, N2^f/f^ and N2^−/−^ mice (8–12 weeks old) were fed a 0.2 % adenine diet (AD) for 28 days. **A**, Representative images of kidney sections stained with Masson’s trichrome, with quantification of fibrotic area (n = 4; scale bar, 100 µm). **B,C**, Blood urea nitrogen (B-UN) and blood creatinine (B-CRE) levels (n = 6–8). **D–F**, Wild-type mice (12 weeks old) were injected with αDll4 (0.02 or 0.1 mg/kg) or αCt (0.1 mg/kg) every 4 days during AD feeding. **D**, Representative Masson’s trichrome-stained kidney sections, with quantification of fibrotic area (n = 4; scale bar, 100 µm). **E,F**, B-UN and B-CRE levels (n = 6). Data are shown as mean ±SEM. P values were determined by two-way ANOVA with Tukey’s post-hoc test (**A–C**), and one-way ANOVA with Tukey’s post-hoc test (**D–F**).

**Fig. S10.**
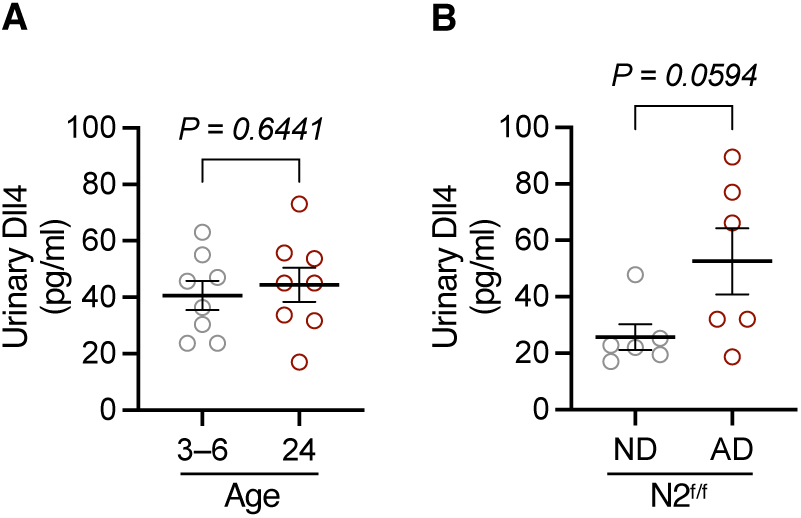
Urinary Dll4 levels are unchanged in aged and CKD mice. A, Urinary Dll4 levels in adult (3–6 months old) and aged (24 months old) mice (n = 8). **B,** Urinary Dll4 levels in N2^f/f^ mice fed a 0.2 % adenine diet (AD) or normal diet (ND) for 28 days. Data are shown as mean ±SEM. P values were determined by Student’s two-tailed unpaired t-test.

**Table S1.** Correlation with serum Dll4 levels.

|  | Spearman's correlation | P-value |
|---|---|---|
| Age | 0.2385 | <0.0001 |
| Height | 0.0339 | 0.1972 |
| Weight | -0.0271 | 0.3026 |
| BMI | -0.0578 | 0.0278 |
| Systolic BP | -0.0753 | 0.0041 |
| Diastolic BP | -0.1686 | <0.0001 |
| Handgrip strength | -0.0323 | 0.2197 |
| Usual gait time | 0.1622 | <0.0001 |
| Maximum gait time | 0.1412 | <0.0001 |
| WBC | 0.0121 | 0.6448 |
| RBC | -0.1340 | <0.0001 |
| Hb | -0.1307 | <0.0001 |
| Hematocrit | -0.1452 | <0.0001 |
| MCV | -0.0009 | 0.9726 |
| MCH | 0.0098 | 0.7084 |
| MCHC | 0.0041 | 0.8758 |
| Platelet | -0.1120 | <0.0001 |
| Total protein | 0.0017 | 0.9472 |
| Albumin | -0.1635 | <0.0001 |
| Total bilirubin | -0.0919 | 0.0005 |
| GOT | 0.1261 | <0.0001 |
| GPT | 0.0012 | 0.9632 |
| ALP | 0.0442 | 0.0924 |
| LDH | 0.0386 | 0.142 |
| γ-GT | -0.0182 | 0.4877 |
| CK | -0.0087 | 0.7394 |
| Total cholesterol | -0.1070 | <0.0001 |
| LDL cholesterol | -0.0573 | 0.0291 |
| HDL cholesterol | -0.1277 | <0.0001 |
| HbA1c | -0.0006 | 0.9827 |
| Glycoalbumin | 0.1286 | <0.0001 |
| BUN | 0.1802 | <0.0001 |
| Creatinine | 0.3622 | <0.0001 |
| eGFR | -0.3952 | <0.0001 |
| Uric acid | 0.2007 | <0.0001 |
| Na | -0.1561 | <0.0001 |
| K | 0.1953 | <0.0001 |
| Cl | 0.0053 | 0.839 |
| TSH | 0.1235 | <0.0001 |
| Free T4 | -0.0372 | 0.1566 |
| HS-CRP | 0.0979 | 0.0002 |
| Urine albumin | 0.1104 | <0.0001 |
| Urine creatinine | 0.0832 | 0.0015 |
| uACR | 0.0633 | 0.0159 |

**Table S2.**
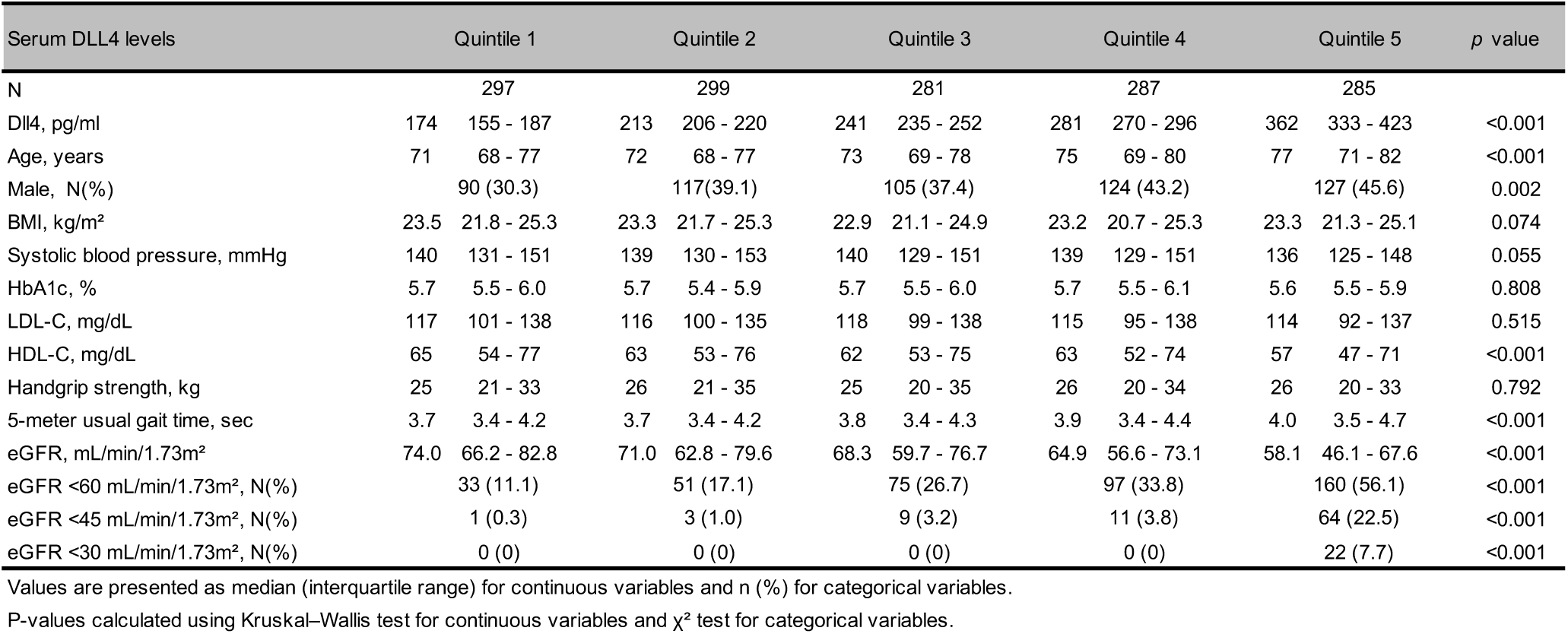
Characteristics of participants according to quintiles of serum Dll4 levels.

| Serum DLL4 levels | Quintile 1 |  | Quintile 2 |  | Quintile 3 |  | Quintile 4 |  | Quintile 5 |  | *p* value |
|---|---|---|---|---|---|---|---|---|---|---|---|
| N | 297 |  | 299 |  | 281 |  | 287 |  | 285 |  |  |
| Dll4, pg/ml | 174 | 155 - 187 | 213 | 206 - 220 | 241 | 235 - 252 | 281 | 270 - 296 | 362 | 333 - 423 | <0.001 |
| Age, years | 71 | 68 - 77 | 72 | 68 - 77 | 73 | 69 - 78 | 75 | 69 - 80 | 77 | 71 - 82 | <0.001 |
| Male, N(%) | 90 (30.3) |  | 117(39.1) |  | 105 (37.4) |  | 124 (43.2) |  | 127 (45.6) |  | 0.002 |
| BMI, kg/m<sup>2</sup> | 23.5 | 21.8 - 25.3 | 23.3 | 21.7 - 25.3 | 22.9 | 21.1 - 24.9 | 23.2 | 20.7 - 25.3 | 23.3 | 21.3 - 25.1 | 0.074 |
| Systolic blood pressure, mmHg | 140 | 131 - 151 | 139 | 130 - 153 | 140 | 129 - 151 | 139 | 129 - 151 | 136 | 125 - 148 | 0.055 |
| HbA1c, % | 5.7 | 5.5 - 6.0 | 5.7 | 5.4 - 5.9 | 5.7 | 5.5 - 6.0 | 5.7 | 5.5 - 6.1 | 5.6 | 5.5 - 5.9 | 0.808 |
| LDL-C, mg/dL | 117 | 101 - 138 | 116 | 100 - 135 | 118 | 99 - 138 | 115 | 95 - 138 | 114 | 92 - 137 | 0.515 |
| HDL-C, mg/dL | 65 | 54 - 77 | 63 | 53 - 76 | 62 | 53 - 75 | 63 | 52 - 74 | 57 | 47 - 71 | <0.001 |
| Handgrip strength, kg | 25 | 21 - 33 | 26 | 21 - 35 | 25 | 20 - 35 | 26 | 20 - 34 | 26 | 20 - 33 | 0.792 |
| 5-meter usual gait time, sec | 3.7 | 3.4 - 4.2 | 3.7 | 3.4 - 4.2 | 3.8 | 3.4 - 4.3 | 3.9 | 3.4 - 4.4 | 4.0 | 3.5 - 4.7 | <0.001 |
| eGFR, mL/min/1.73m<sup>2</sup> | 74.0 | 66.2 - 82.8 | 71.0 | 62.8 - 79.6 | 68.3 | 59.7 - 76.7 | 64.9 | 56.6 - 73.1 | 58.1 | 46.1 - 67.6 | <0.001 |
| eGFR <60 mL/min/1.73m<sup>2</sup>, N(%) | 33 (11.1) |  | 51 (17.1) |  | 75 (26.7) |  | 97 (33.8) |  | 160 (56.1) |  | <0.001 |
| eGFR <45 mL/min/1.73m<sup>2</sup>, N(%) | 1 (0.3) |  | 3 (1.0) |  | 9 (3.2) |  | 11 (3.8) |  | 64 (22.5) |  | <0.001 |
| eGFR <30 mL/min/1.73m<sup>2</sup>, N(%) | 0 (0) |  | 0 (0) |  | 0 (0) |  | 0 (0) |  | 22 (7.7) |  | <0.001 |

Values are presented as median (interquartile range) for continuous variables and n (%) for categorical variables.
P-values calculated using Kruskal–Wallis test for continuous variables and χ<sup>2</sup> test for categorical variables.

**Table S3.** Odds ratios for low estimated glomerular filtration rate (eGFR) across quintiles of serum Dll4 levels.

| Dll4 Category | Model 1(Crude) |  |  |  | Model 2 |  |  |  | Model 3 |  |  |  | Model 4 |  |  |  |
|---|---|---|---|---|---|---|---|---|---|---|---|---|---|---|---|---|
|  | OR (95% CI) |  |  |  | OR (95% CI) |  |  |  | OR (95% CI) |  |  |  | OR (95% CI) |  |  |  |
| Q1 (lowest) | 1 | (reference) |  |  | 1 | (reference) |  |  | 1 | (reference) |  |  | 1 | (reference) |  |  |
| Q2 | 1.65 | 1.03 | - | 2.63 | 1.61 | 1.00 | - | 2.60 | 1.63 | 1.01 | - | 2.65 | 1.63 | 1.00 | - | 2.65 |
| Q3 | 2.91 | 1.86 | - | 4.56 | 2.83 | 1.79 | - | 4.47 | 2.99 | 1.88 | - | 4.76 | 2.99 | 1.88 | - | 4.76 |
| Q4 | 4.08 | 2.64 | - | 6.32 | 3.46 | 2.21 | - | 5.41 | 3.73 | 2.37 | - | 5.86 | 3.73 | 2.37 | - | 5.86 |
| Q5 (highest) | 10.24 | 6.66 | - | 15.76 | 8.12 | 5.21 | - | 12.66 | 8.69 | 5.54 | - | 13.65 | 8.73 | 5.55 | - | 13.72 |
| p for trend | <0.001 |  |  |  | <0.001 |  |  |  | <0.001 |  |  |  | <0.001 |  |  |  |

Model 1: Unadjusted.
Model 2: Adjusted for age and sex.
Model 3: Model 2 + body mass index, systolic blood pressure,  HbA1c, LDL cholesterol, HDL cholesterol.
Model 4: Model 3 + grip strength and 5-meter usual gait time.

**Table S4.**
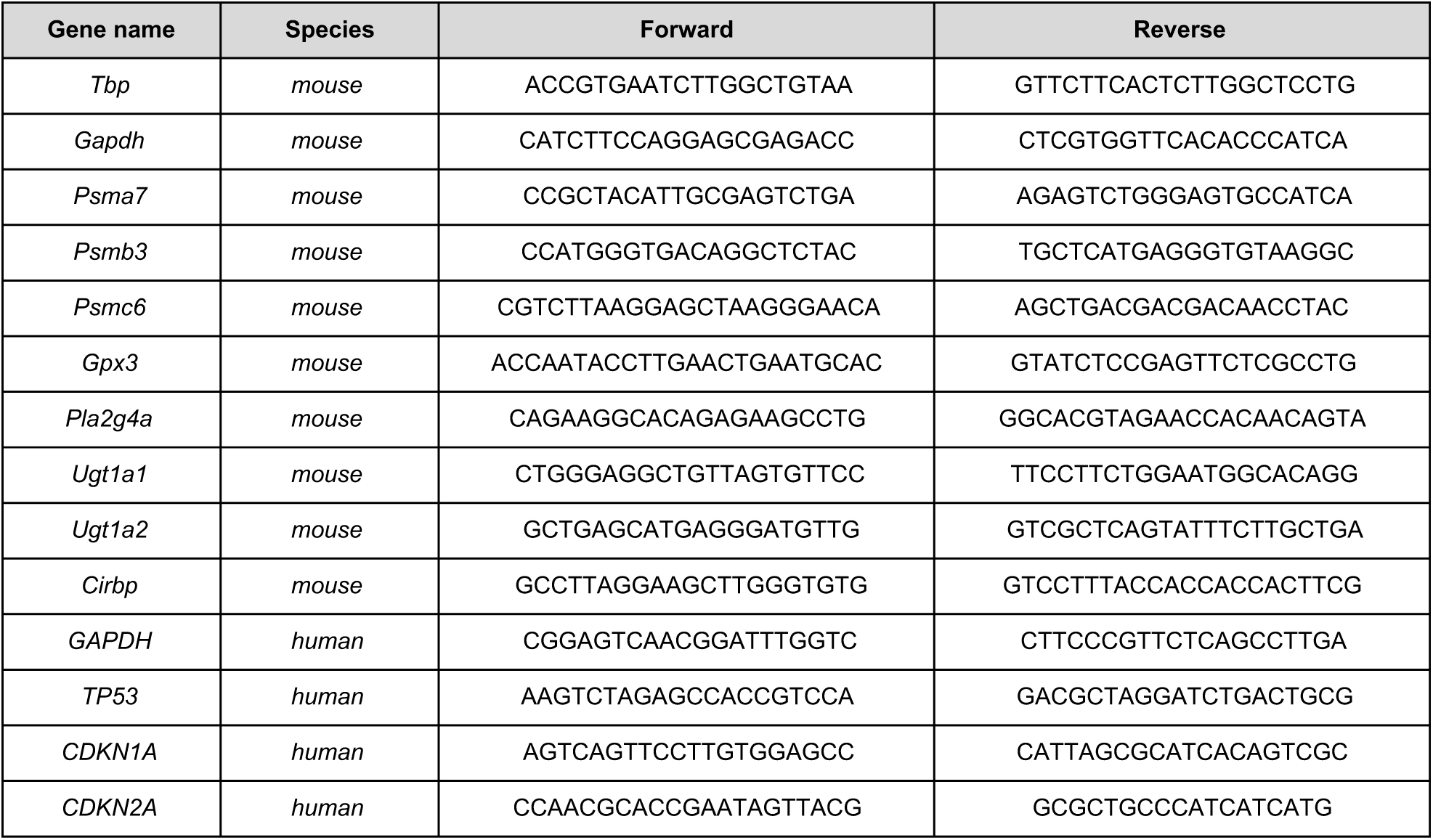
List of primers used in this study.

